# Complex Human hear bearing Skin Organoids reveal Cell Type Specific Susceptibility and Innate Immune Responses to Herpes Simplex Virus 1

**DOI:** 10.1101/2025.02.10.637415

**Authors:** Emanuel Wyler, Silvia Albertini, Caroline C. Friedel, Artür Manukyan, Annalena Winterfeldt, Nina Bracker, Jiabin Huang, Susanne Krasemann, Izabela Plumbom, Janine Altmüller, Thomas Conrad, Rudolph Reimer, Carola Schneider, Helena Radbruch, Wali Hafezi, Arne Hansen, Adam Grundhoff, Markus Landthaler, Nicole Fischer, Manja Czech-Sioli

**Affiliations:** Max Delbrück Center for Molecular Medicine in the Helmholtz Association (MDC), Berlin Institute for Medical Systems Biology, Berlin, Germany; Institute for Molecular Virology and Tumorvirology, University Medical Center Hamburg-Eppendorf Hamburg, Germany; Institute for Informatics, Ludwig-Maximilians-Universität München, Munich, Germany; Institute of Medical Microbiology, Virology and Hygiene, University Medical Center Hamburg-Eppendorf Hamburg, Germany; Institute of Neuropathology, University Medical Center Hamburg-Eppendorf, Hamburg, Germany; Genomics Technology Platform, Max Delbrück Center for Molecular Medicine in the Helmholtz Association, Berlin, Germany; Core Facility Genomics, Berlin Institute of Health at Charité - Universitätsmedizin Berlin, Berlin, Germany; Technology platform Microscopy and Image Analysis/Electron Microscopy, Leibniz Institute of Virology, Hamburg, Germany; Department of Neuropathology, Charité – Universitätsmedizin Berlin, Berlin, Germany; University Hospital Muenster, Institute of Virology / Clinical Virology, Muenster, Germany; Institute of Experimental Pharmacology and Toxicology, University Medical Center Hamburg-Eppendorf, Hamburg, Germany; Department of Viral Genomics, Leibniz Institute of Virology, Hamburg, Germany; Institut für Biologie, Humboldt-Universität zu Berlin, Berlin, Germany

## Abstract

Epithelial surfaces are the initial site of Herpes Simplex Virus 1 (HSV-1) infection. After latency establishment in the peripheral nervous system, reactivation can lead to skin pathologies. To reflect the complexity of skin we used complex human hair bearing skin organoids as a new model system for HSV-1 infection. We analyzed spread of the infection through the three-dimensional tissue and the cell type specific host response in a spatial temporal manner. Single-cell-resolution spatial transcriptomics and imaging shows a restricted viral infection in keratinocytes of the epidermis and hair follicles. The host factor IER3 is upregulated in papillary fibroblasts of the dermis as the infection progresses through the tissue. In keratinocytes of the stratum basale, we find IRF3 upregulated in infected but also virus negative cells, suggesting a signaling mechanism to initiate antiviral responses. Furthermore, we find a cell type specific inflammatory response with an upregulation of TNFSF9 in dermal fibroblasts and TNF in keratinocytes. TNFα target genes are induced in neighboring uninfected cells suggesting the induction of a localized inflammatory reaction to suppress viral infection. In summary, our findings position human hair-bearing skin organoids as a highly physiological model system for studying HSV-1 pathogenesis and reveal spatially organized host responses.

## Introduction

Herpes Simplex Virus 1 (HSV-1) is a highly prevalent human pathogen with a global prevalence of ∼67% in under 50-year-old individuals^1^. Primary infection usually occurs via the mucosal surfaces or micro-lesions in the skin. Initially HSV-1 productively replicates in the epithelial cells of skin and mucosa causing inflammation and tissue damage and consequently blisters^2^. After replication in epithelial cells, HSV-1 enters the nerve endings of peripheral neurons and travels along their axons to the cell bodies in trigeminal ganglia where it establishes latency^3,4^. Upon spontaneous reactivation, HSV-1 enters the lytic life cycle and produces infectious viral particles that migrate in an anterograde manner to the epithelial cells of skin and mucosa. The following lytic replication in the epithelium leads to the characteristic herpes lesions that release virus to the environment^2^.

In the past, many different model systems have been used to elucidate the mechanisms of primary infection and reactivation by HSV-1 in the skin. Due to the complexity of skin and the involvement of many different cell types, *in vitro* modelling by monolayer cell cultures can only recapitulate certain aspects of the viral life cycle. More complex model systems used to study virus susceptibility, spread and cytopathic effects are animal models^5–8^, *ex vivo* murine^8–10^ and human skin explants^8,11,12^, and epithelial raft cultures^13,14^.

While the understanding of HSV-1 pathogenesis in the skin has exceptionally increased using these model systems, many aspects remain elusive. Especially the susceptibility and contributions of specific structures, like hair follicles, different fibroblasts subpopulations and low abundancy cell types, to primary infection, latency establishment and reactivation are not understood. The field of organoids is constantly evolving and creating potential new model systems for virus research^15–17^. In this study we used a highly complex skin organoid (SkO) model generated from human induced pluripotent stem cells (hiPSCs) for the analysis of HSV-1 pathogenesis in the skin^18,19^. SkOs contain a stratified epidermis, fat-rich dermis, hair follicles (HFs) producing pigmented hair, sebaceous glands, melanocytes and Merkel cells. The HFs are innervated by sensory neurons that are covered in Schwann cells on their axons while their somata assemble in trigeminal like clusters together with satellite glial cells. SkOs contain all cell types expected in the fetal skin of the second trimester of development except for sweat glands, vascularization and immune cells^18,19^. Similar to many other organoids, SkOs show an inside out morphology with the dermis being exterior and exposed to the culture medium and the epidermis being interior with hair from hair follicles growing to the inside of the organoid. In this study, SkOs were infected with HSV-1 from the dermal side, leading to viral spread from the dermis to the epidermis, thereby recapitulating primary infection associated with deep dermal wounding or viral reactivation. We analyzed the susceptibility of different cell types to HSV-1 infection and found a restricted HSV-1 infection in keratinocytes as well as in HF cells of both dermal and epidermal origin. Furthermore, we found the host factors IER3 and IRF3 upregulated in a spatial temporal manner in response to the infection. We found a spatially restricted inflammatory response by expression of TNF and TNFSF9 leading to downstream activation of TNF target genes. Overall, our study shows how complex infection models combined with high-resolution spatial omics methods allow the investigation of cellular responses and the progression of infection through complex three-dimensional tissues.

## Results

### Infection of skin organoids (SkOs) with HSV-1

SkOs were produced after the protocol from Lee and collegues^19^ and checked for correct differentiation by brightfield microscopy, immunohistochemistry (IHC) and whole-mount immunofluorescence (WMI) staining (Supplementary Figure 1A-D). We observed the expected full complexity at ∼120 days of differentiation with the dermis and epidermis in the head structure and cartilage in the tail. The keratinocytes in the epidermis showed a basal layer and differentiation into stratum spinosum and granulosum with terminally differentiated dying keratinocytes accumulating in the head of the SkO (Supplementary Figure 1D). Hair follicles (HFs) showed the expected structures i.e., dermal papilla, matrix, inner root sheath, and outer root sheath. We detected Merkel cells and innervation of the HFs with assembly of neuronal somata in the organoid tail (Supplementary Figure 1B-D).

SkOs at 120 days of differentiation were infected with HSV-1 strain 17 expressing GFP under the control of a CMV IE-promoter. To observe spread of the infection within the tissue from single spots, infection was performed with a low viral dose (800 PFU per SkO). At 2dpi GFP expression was visible by fluorescence microscopy at spots on the surface of the organoid (Figure 1A). In IHC staining the first layers of fibroblasts in the dermis showed expression of GFP and HSV-1 glycoprotein D (gD) and indications of a cytopathic effect (CPE). Infection spread solely among cells of the dermis with exclusion of the HFs (Figure 1B, Supplementary Figure 2). At 4dpi GFP fluorescence increased and covered the entire organoid (Figure 1A). IHC staining for GFP and gD showed that the infection reached the inner structures of the organoid including the HFs and keratinocytes of the basal layer and the stratum spinosum.

**Figure 1:**
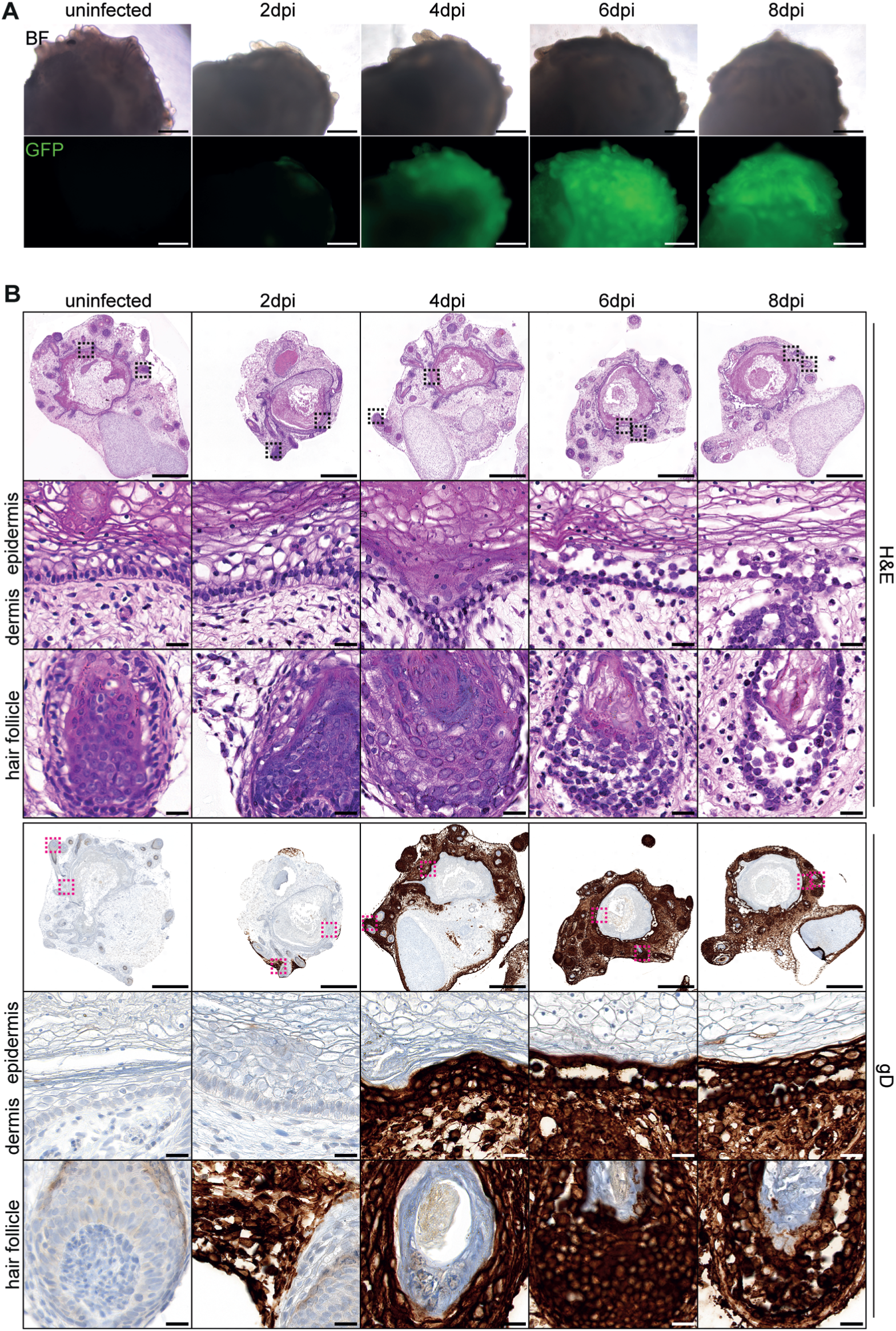
Infection of skin organoids with HSV-1 leads to cytopathic effects. **A:** Brightfield (BF) and fluorescence (GFP) images of skin organoids (SkOs) infected with HSV-1 (expressing GFP from a CMV-IE promoter) at 120 days of differentiation. Shown are uninfected and HSV-1 infected SkOs at 2, 4, 6, and 8 days post infection (dpi). Scale bars 500µm. **B:** IHC staining of SkOs infected with HSV-1 as shown in A. Boxes in overview images indicate areas of higher magnification shown below. Scale bars 500µm in overview images and 20µm in zoom-ins. Upper picture of zoom-ins shows epidermis on the top and dermis below. Lower picture in zoom-ins shows hair follicle. H&E: Hematoxylin and eosin stain. gD: Staining against HSV-1 glycoprotein D.

Terminally differentiated keratinocytes showed no signs of infection. Similarly, the different cell layers at the HFs were positive for GFP and gD except for the cornified hair structures (Figure 1B, Supplementary Figure 2). Until 8dpi, we observed no sign of infection in these structures, as in the cartilage of the organoid tail. At 4dpi a CPE could be observed mainly in the dermis. At 6dpi infected keratinocytes in the epidermis and the inner cell layers at the HFs started to show a CPE that resulted in dissolvement of the basal membrane and HF integrity. Taken together, these findings showed that HSV-1 infects almost all cell types of SkOs with indications of differential spreading in different cell types.

### Bulk transcriptomics of HSV-1 infected skin organoids

To analyze the dynamics of viral gene expression and the host response to the viral infection we performed bulk RNA-seq of uninfected and HSV-1 infected SkOs at 2, 4, 6 and 8dpi. At 2dpi we observed only 20 differentially expressed genes (DEGs), which increased to ∼1000 at 4dpi and ∼5000 at 6 and 8dpi (Figure 2A, Supplementary Figure 2B, C). At 4dpi vs. uninfected we found an enrichment of GO-terms of inflammation (e.g., NFκB signaling, response to tumor necrosis factor) in the upregulated differentially expressed genes (Figure 2B). At 6 and 8dpi we found terms of cellular differentiation (e.g., skin development, molting cycle) stand out among the downregulated DEGs. This could reflect the typical HSV-1 induced host cell shutoff^20^ and the observed CPE. However, the percentage of viral reads reaches ∼20% of all reads already at 4dpi and increases only slightly up to ∼25% at 6 and 8dpi (Figure 2C). Overall, this data indicated that key transcriptional changes are to be expected during the first 4 to 6dpi, setting the time frame for subsequent experiments.

**Figure 2:**
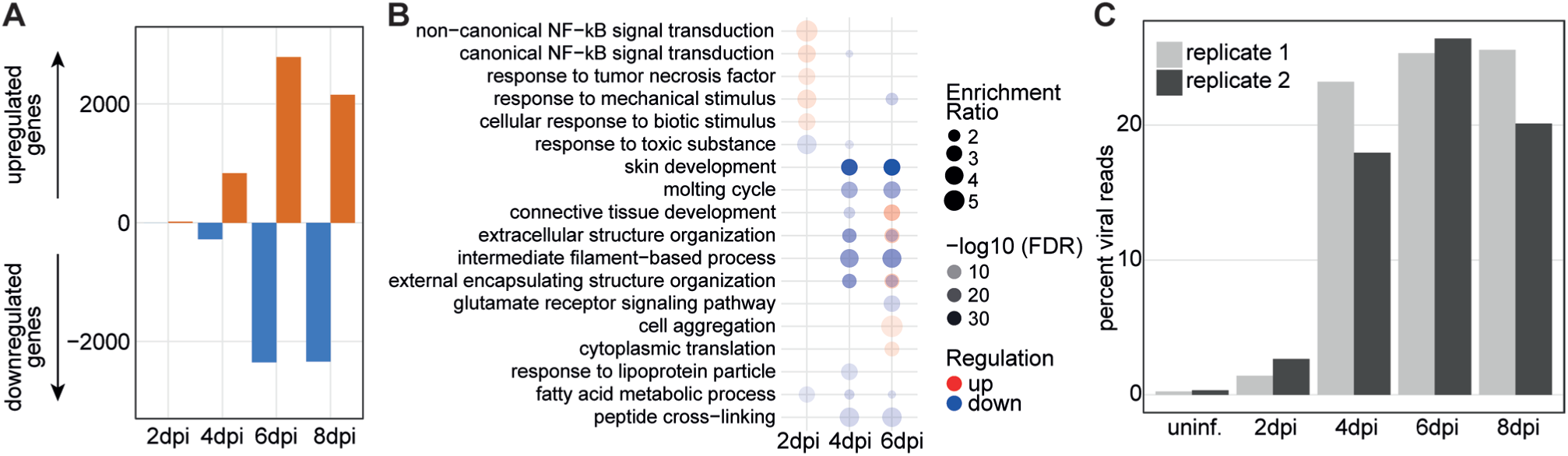
Bulk RNA-seq of HSV-1 infected SkOs. **A:** Barplot showing numbers of differentially expressed host genes at 2, 4, 6, and 8dpi versus uninfected SkOs. **B:** GO analysis (WebGestalt) was performed using significantly (padj. ≤0.05) up- or downregulated genes (log2FC ≥1/≤-1) from RNA-seq analysis of SkOs infected with HSV-1 at 4, 6 and 8dpi compared to uninfected SkOs. Shown are the top 5 significantly enriched GO-Terms (FDR ≤0.01, sorted by enrichment ratio) in the up- and downregulated genes. At 2dpi no GO-terms were significantly enriched. At 4dpi only 2 GO-terms were associated with the downregulated genes. At 6dpi no GO-terms were significantly enriched in the upregulated genes. **C:** Percentage of total reads mapping to HSV-1 in uninfected and 2, 4, 6, 8dpi. Each bar shows one replicate.

### Spatial transcriptomics of HSV-1 infected skin organoids

To understand the temporal spatial dynamics of the viral infection and the host response in the SkO model system we performed two spatial transcriptomics experiments with single cell resolution on the Xenium platform. In the pilot experiment, we profiled sections of each 3 SkOs uninfected, at 2dpi, and 6dpi. Due to a strong CPE and HSV-1 induced host cell shutoff, we were not able to perform cell type annotation for most cells at the 6dpi time point (see Supplementary Note for results of the pilot experiment). Therefor, we switched to 2dpi, 3dpi and 4dpi timepoints in the main experiment (Figure 3, Supplementary Figure 3). We quantified the expression of the 470 measured genes, including five viral transcripts (US1, UL27, UL29, UL54 and LAT). Cell type identities were assigned based on cellular transcriptomes (Figure 3A-D, Supplementary Figure 3A,C,D, see Supplementary Note for cell type assignment strategy). We identified the expected cell types based on single-cell RNA-sequencing data and immunostainings performed by Lee and colleagues^18,21^. Additionally, we could distinguish between subtypes that coincide with specific localization such as different fibroblast types (e.g., papillary 1, 2 and mesenchymal) that localize proximal or distal to the epidermis. Keratinocyte subtypes at different stages of differentiation were distinguishable and clearly spatially restricted. We annotated the stratum basale and the stratum corneum while the stratum spinosum and granulosum were combined as one cluster, since the markers in our panel were not sufficient for separation. Cell types of the HFs (dermal sheath, outer root sheath, inner root sheath, matrix, bulge region, dermal papilla) were clearly identified and matched their expected localization. Cell type assignments were also successful in infected SkOs (Figure 3A, D), despite downregulation of host transcripts in infected cells. This can be visualized e.g., with complete loss of PDGFRα mRNA, a pan-fibroblast marker, in highly infected cells due to host cell shutoff^20^ (Figure 3B, top panels). However, some highly infected cells, particularly fibroblasts, lost their transcriptional identity and were thus assigned as “undefined” (Figure 3A, gray colored cells, Supplementary Figure 3A). Accordingly, the proportion of “undefined” cells increased in highly infected (4dpi) vs. uninfected organoids but did not exceed ∼15% of all cells (Figure 3D).

**Figure 3:**
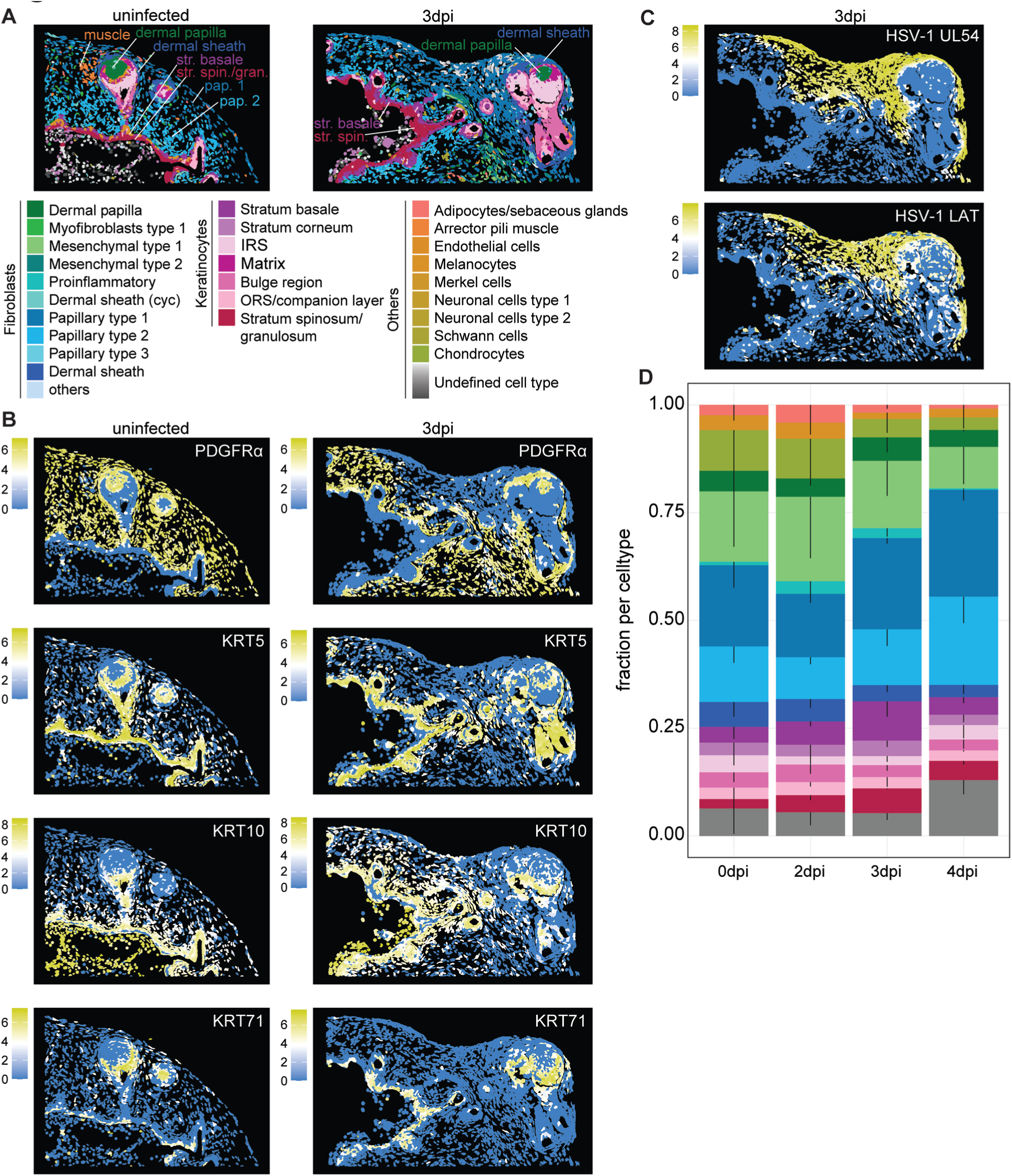
Spatial transcriptomics analysis of HSV-1 infected SkOs. **A:** Representative regions from an uninfected SkO (left) and a SkO at 3dpi (right), with cells colored by annotated cell type. **B, C:** The same organoid regions as in A, colored by expression levels of the indicated genes. **D:** Relative abundances of major cell types across the indicated timepoints. Thin lines denote standard deviations calculated across three organoids per time point. Cell type color coding is consistent with panel A.

Analysis of the expression of the viral transcripts US1/ICP22, UL54/ICP27, UL29/ICP8, UL27/gB and LAT matched the protein staining, showing infection starting in the outermost fibroblast layers of the dermis at 2dpi and then spread through the organoid until the infection reached the epidermis at 4dpi (Figure 3C and Supplementary Figure 3B).

### Susceptibility to viral infection varies widely between cell types

To investigate susceptibility and viral gene expression between different cell types in SkOs, we considered the special features of an infection spreading in a tissue compared to an infection in a monolayer cell culture. HSV-1 infects the outer layers of the organoids first and later the more inner parts leading to an asynchronous infection. First, we calculated the proportion of infected cells and the viral load in the most abundant subtypes, papillary fibroblasts type 1/2, keratinocytes of the stratum basale and stratum spinosum/granulosum, and four cell types in HFs (dermal sheath, dermal papilla, outer root sheath, inner root sheath; Figure 4A). At 3dpi papillary fibroblasts type 1, which is the outermost cell layer of SkOs, and the dermal sheath fibroblasts had the highest number of HSV-1 infected cells (∼25%). Also, papillary fibroblasts type 1 had a significantly higher viral load than all other cell types including the dermal sheath. At 4dpi, when the infection reaches the inner parts of the organoid, the proportion of infected cells increased in all cell types. However, stratum basale and spinosum/granulosum keratinocytes still showed significantly less infected cells than papillary fibroblasts (Figure 4A, upper panel). Viral loads were significantly lower than in papillary fibroblasts type 1 in all cell types except papillary fibroblasts type 2 (Figure 4A, lower panel). To exclude that this effect was caused by the asynchronous infection spreading from the outside to the inside we compared stratum basale keratinocytes with papillary fibroblasts at equal distance ranges from the exterior surface (Supplementary Figure 4AB). Again, we found significantly lower viral loads. Furthermore, we analyzed the influence of infectious cells in proximity to basal keratinocytes (<50µm distance) (Supplementary Figure 5A). Infectious cells were defined as cells with 20% or more viral transcripts. As expected, we found an increase of the viral load with rising numbers of infectious cells in the neighborhood. Interestingly, papillary fibroblasts reached their maximum viral load (up to 70% viral transcripts) with 1 to 2 infectious cells in proximity. Keratinocytes of the stratum basale showed a significantly lower viral load compared to papillary fibroblasts despite the same number of infectious cells in their neighborhood. Even with >6 infectious cells in proximity the mean viral load did not exceed 25% of all transcripts. To analyze the dynamics of the viral life cycle we ordered and binned infected cells along increasing viral load. We found in papillary fibroblasts a strong increase of UL54 RNA at the beginning followed by a plateau, with the LAT RNA constantly accumulating (Figure 4B, top panel). In stratum basale keratinocytes, there was an indication of a longer lag phase, and a slower increase of viral gene expression (Figure 4B, bottom panel). These results suggest that keratinocytes of the stratum basale have a delayed or restricted viral life cycle. In dermal sheath fibroblasts, the dermal papilla and the keratinocyte subtypes of HFs (e.g., matrix, root sheaths) we also observed significantly lower viral loads than in the surrounding papillary fibroblasts (Figure 4A). Staining for the viral early protein ICP8 and the late protein gC showed that viral infection spreads first into the dermis and excludes the HF layers (Figure 4C). HF cell layers are infected after the infection already spread to the inner parts of the organoids at 4dpi with the dermal papilla being infected last (Figure 4C, right panel) but eventually infection reaches all layers of HFs with a CPE being observable at 6 to 8dpi (Figure 1B). We hypothesized three potential explanations for this observation. First, there could be a layer of non-permissive cells surrounding the HFs. Second, a barrier of extracellular matrix might act protectively. And third, the infection cycle in the different cell types within the HFs could be slower compared to e.g., the surrounding papillary fibroblasts. To investigate these possibilities, we defined dermal sheath fibroblasts located next to highly infected papillary fibroblasts (Supplementary Figure 5B). Indeed, the viral load in these dermal sheath fibroblasts showed a strong drop from the cells outside in direct proximity (Figure 4E). As for keratinocytes of the stratum basale, the lower viral load in cell types of HFs was not driven by spatial effects like the localization relative to the organoid boundary or a lower number of infected cells in proximity (Supplementary Figure 4AB, Supplementary Figure 5A). IF staining of the analyzed cells showed close contact between dermal sheath and papillary fibroblasts with no indication of any physical barriers between these cell layers (Supplementary Figure 5C). This is supported by electron microscopy which showed close proximity of papillary fibroblasts and dermal sheath fibroblasts (Figure 4F, Supplementary Figure 6). We observed that at 3dpi many viral particles are present in the papillary fibroblasts layer inside cells (nucleus/cytoplasm) as well as in the extracellular space (Figure 4F, zoom-ins). In the dermal sheath layer, the matrix, the outer, and inner root sheath we found only a small number of viral particles in the cytoplasm and the nucleus but not in the extracellular space. In the nuclei we found viral A and C capsids indicating viral replication (A-capsids lack DNA, B capsids lack DNA but contain the scaffolding protein and C-capsids contain the viral DNA^22^) (Figure 4F, 3dpi, zoom-in on matrix). In the inner layers of the HF we did not find any viral particles. However, at 4dpi we find viral particles in the extracellular space, the cytoplasm, and the nucleus in almost all HF layers present in our sections (dermal sheath, outer root sheath, matrix, inner root sheath cuticle, companion layer, Huxley’s layer, dermal papilla) except for the Henle’s layer (Figure 4F, Supplementary Figure 6B). Again, we found A- and C- but also B-capsids in replication compartments of the nucleus in these HF cell types. From these results we conclude that HSV-1 lytically replicates in most cell types of HFs but similar to keratinocytes of the stratum basale viral replication is delayed in these cells which slows down viral spread.

**Figure 4:**
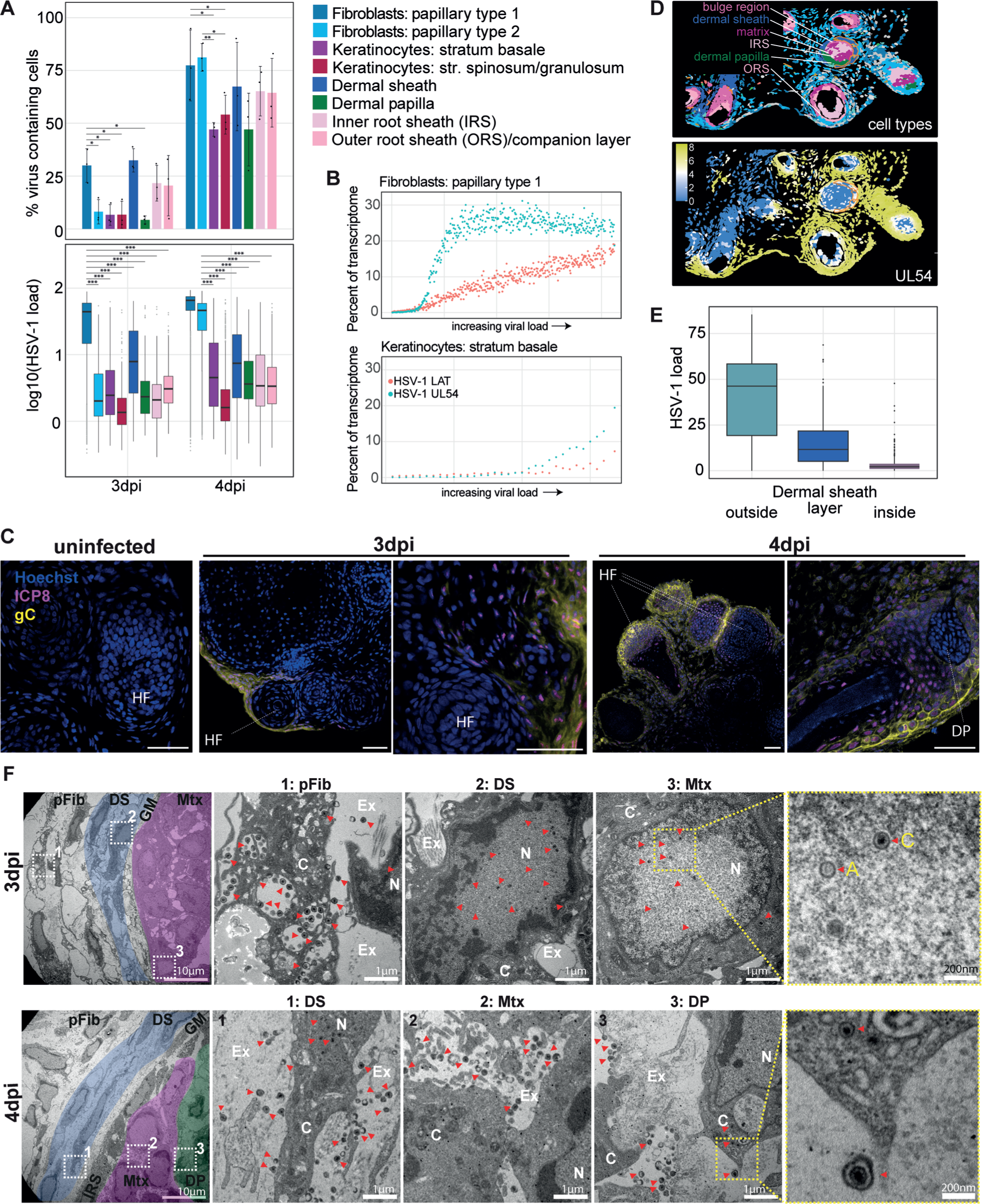
Susceptibility to viral infection varies widely between cell types. **A:** Upper part: percent of cells with at least 2 HSV-1 transcript counts for selected cell types. Indicated are individual values from the three organoids per timepoint, and standard deviations. * p-value < 0.05 ** p-value < 0.01 (Student’s t-test). Lower part: distribution of log10-transformed viral loads (percent viral transcripts). The middle line in the boxplot displays the median, the box indicates the first and third quartile, whiskers the 1.5 interquartile range (IQR). Outliers beyond are marked by single dots. *** p-value < 0.001 (Wilcoxon rank sum test). **B:** Cells with at least 2 viral counts were ordered according to viral load and binned with 20 cells per bin. Shown are the percentages of the two indicated viral transcripts within the bin for papillary fibroblasts type 1 (top) and stratum basale keratinocytes (bottom). **C:** Sections from WMI staining of HSV-1 infected SkOs harvested at the indicated dpi and stained for HSV-1 proteins ICP8 and gC. Scale bars: 50µm. **D:** HSV-1 UL54 expression around HFs. Shown is a region from a SkO at 4dpi. Upper part: cells colored by cell type, with hair follicle cell types labeled. Lower part: cells colored by normalized HSV-1 UL54 expression. Surrounded with orange are dermal sheaths cells in proximity to highly infected papillary fibroblasts. **E:** Box plot of HSV-1 load (percent viral transcripts) in HFs cells. Dermal sheath layer: Dermal sheath cells next to highly infected cells (surrounded with orange in D). Outside/ inside: other cells within 50µm distance from dermal sheath layer outside/ inside of HFs. **F:** Transmission electron microscopy (TEM) of HFs from HSV-1 infected SkOs at 3dpi and 4dpi. Left images show overviews with colored layers of HFs. pFib: papillary fibroblasts; DS: dermal sheath; GM: glassy membrane; Mtx: matrix; IRS: inner root sheath; DP: dermal papilla; Ex: extracellular space; N: nucleus; C: cytoplasm. Boxes with white and yellow lines, respectively indicate areas of zoom-ins. Red arrows indicate viral particles. A and C capsids are indicated in a zoom-in on a nucleus of a matrix cell at 3dpi.

### IER3 is induced at the infection front

HSV-1 infection leads to a general shutoff of host genome transcription ^20^ as well as read-through ^23^ and antisense transcription ^24^, but also induction of several genes ^25–28^. We used the spatial transcriptomics data to investigate how host gene expression changes are related to the spread of the virus through the tissue. First, we focused on papillary fibroblasts, the most abundant infected cell type. We subclustered cells based on a set of 190 genes deregulated upon infection in papillary fibroblasts (Figure 5AB). By projecting the clusters back onto the section of a 3dpi organoid shown in Figure 3, we observed that the transcriptome-based clustering is reflected in space (Figure 5C). On the one hand, this allowed us to estimate the initial infection site from which the virus gradually spread through the tissue. On the other hand, we could use the gene expression data to investigate what happens at, before and after the “infection front” in a spatial-temporal manner. We calculated DEGs between sets of papillary fibroblasts at different locations (Supplementary Figure 7A-C) which showed that the host gene IER3 is strongly induced at the infection front, with subsequently decreasing in cells later in the viral life cycle (Figure 5C). This is consistent with the transient changes observed in bulk RNA-seq (Supplementary Figure 7D). IER3 is a target gene of the MAP kinase pathway^29^ and HSV-1 infection has been shown to stabilize IER3 through ICP27 via this pathway^30–32^. To corroborate our finding, we performed IER3/ICP8 co-immunostaining in whole-mount organoids (Figure 5D). Also, the IER3 protein appears immediately upon infection, but in contrast to the mRNA, it was also upregulated in cells after the infection front that are later in the viral life cycle. Overall, this example shows how genes can be transiently upregulated at the infection front as infection progresses through the tissue, thus creating a specific virus-host cell interaction microenvironment.

**Figure 5:**
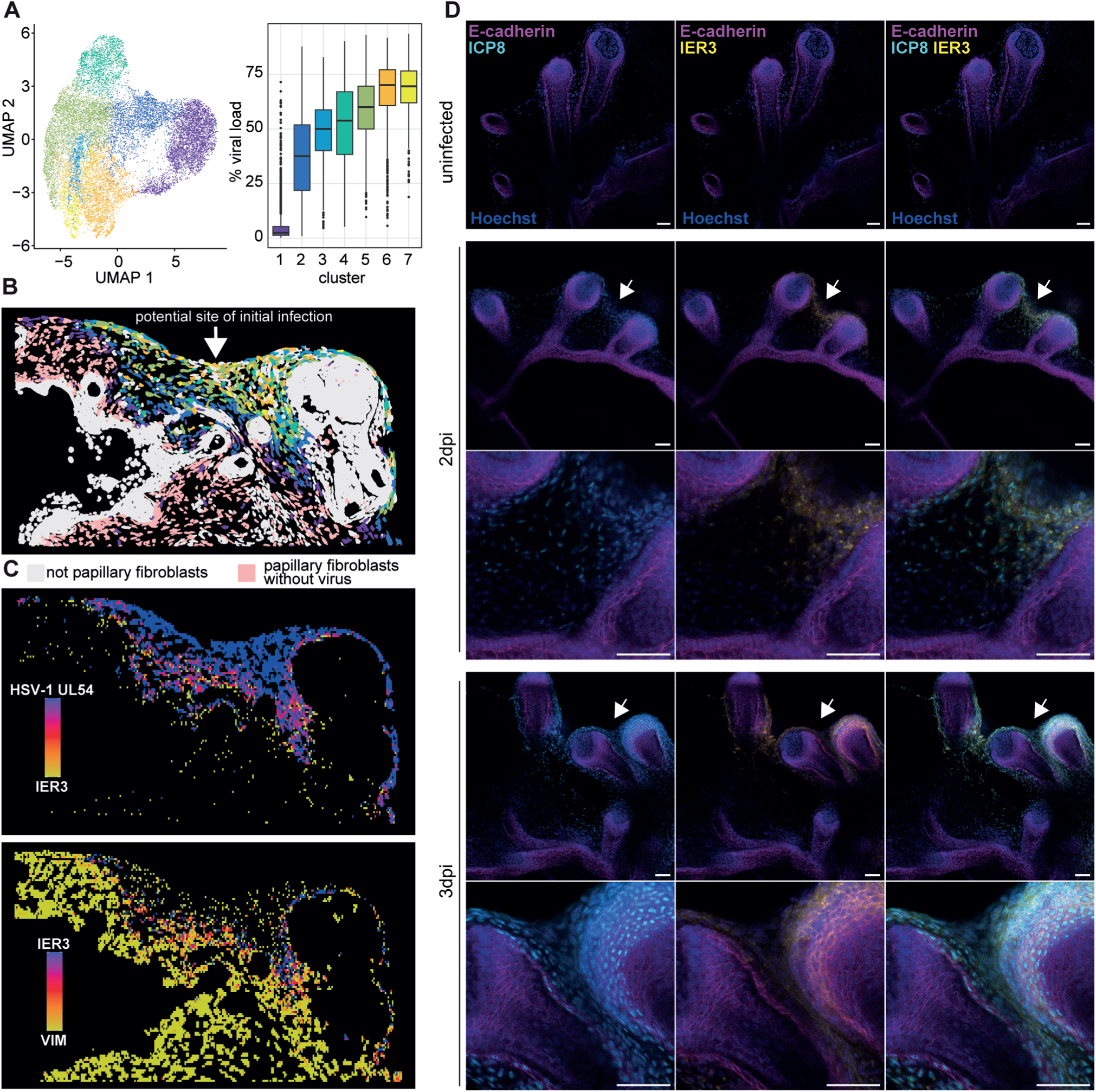
IER3 is induced at the HSV-1 infection front in SkOs. **A:** Papillary fibroblast types 1 and 2 with at least two viral RNA counts were clustered based on viral gene expression and infection-induced host gene dysregulation, resulting in seven distinct clusters. Left: Two-dimensional projection of cells colored by cluster membership. Right: Boxplot showing the percentage viral transcripts in the cells of each cluster. **B:** Region of an infected organoid at 3dpi shown in Figure 3. Papillary fibroblasts are colored by cluster membership as in A, or in light pink if containing less than two viral RNA counts (papillary fibroblasts without virus). Other cell types, including “undefined” cells which includes highly infected papillary fibroblasts are in light grey. **C:** Shown are transcript densities in papillary fibroblasts for genes as indicated in spatial bins. Red colors indicate that both transcripts are present. VIM: Vimentin, general marker for fibroblasts. **D:** Sections from WMI of uninfected and HSV-1 infected SkOs at 2dpi and 3dpi stained for ICP8, IER3 and E-cadherin (marker for epidermis). Arrows indicate potential site of viral infection that is shown in higher magnification below. Scale bars: 50µm.

### HSV-1 infection upregulates the transcription factor IRF3 in keratinocytes

Since defining an “infection front” requires a high number of infected cells from a specific cell type, we could not perform this approach with the lower abundant cell types in our model system. To identify here host genes responding to the infection, we calculated differential expression values per cell type between infected and uninfected organoids. To take the viral RNA content into account, we separated cells into (i) no viral RNA (less than 2 HSV-1 counts), (ii) low virus RNA (viral load below the median calculated within the cell type and time point but across all individual organoids) and (iii) high virus RNA (viral load above the median) (Figure 6A). As positive controls, DUX4, antisense and read-through transcripts were all found to be induced by HSV-1 infection, although to different extent in different cell types. (Supplementary Figure 8A).

**Figure 6:**
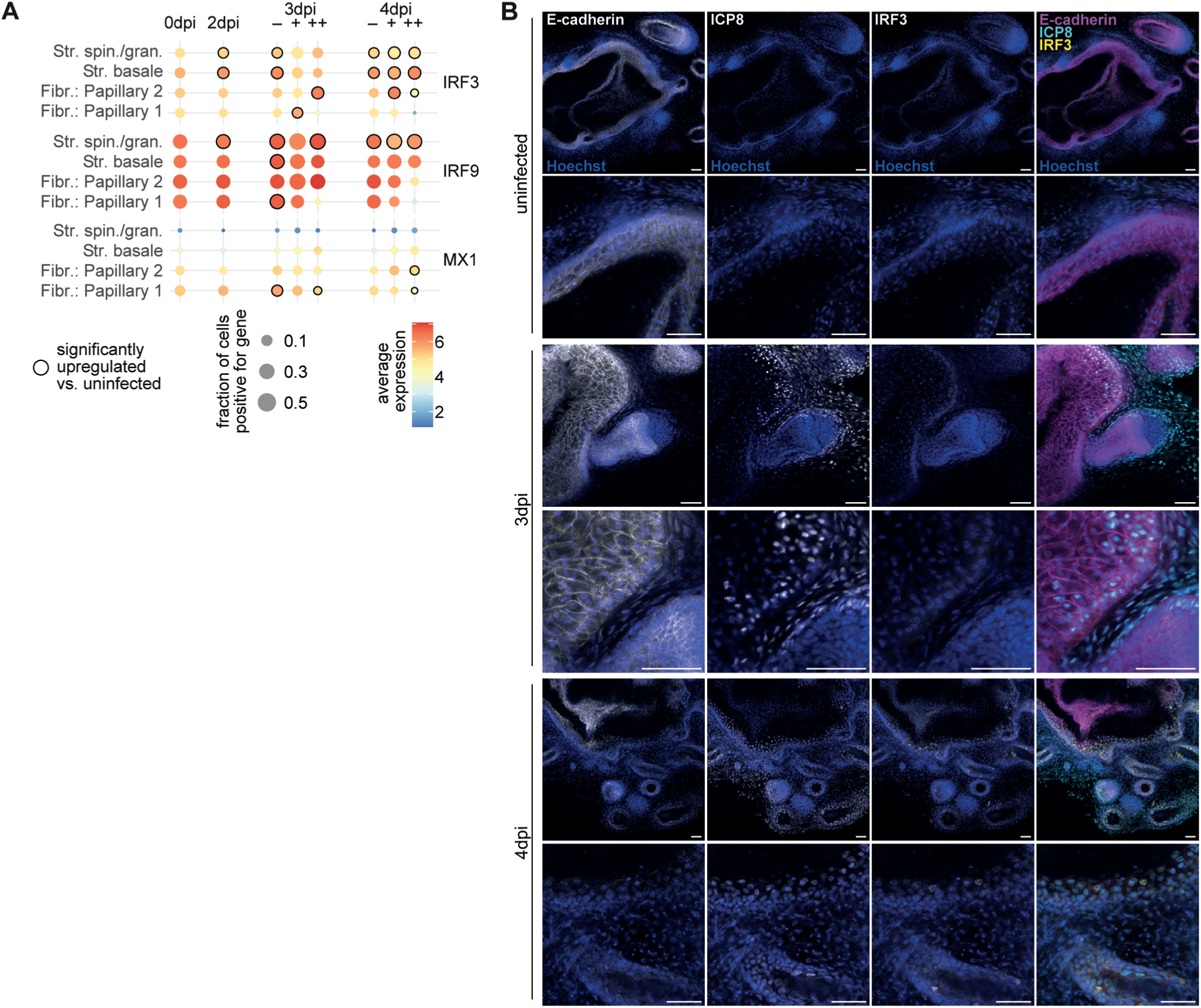
The transcription factor IRF3 is induced in keratinocytes. **A:** Expression values of the indicated genes in a subset of cell types. For uninfected and 2dpi SkOs, all cells were combined for the analysis. For 3dpi and 4dpi, cells are split up in those with no or only 1 viral count (-), and those with viral load below (+) and above (++) the median. The median viral load was calculated only within cells with more than 1 count. Dot sizes represent the fraction of cells having at least one count of the specified gene, the color the average expression. Black circles around the dots indicate significant differential expression in a pseudobulk comparison vs. all cells in uninfected cells (padj <0.05). **B:** Sections from WMI of uninfected and HSV-1 infected SkOs at 3dpi and 4dpi stained for ICP8, IRF3 and E-cadherin (marker for epidermis). Scale bars: 50µm.

For further analysis, we focused on genes involved in interferon signaling and present in the gene panel, namely the transcription factors interferon regulatory factor 3 and 9 (IRF3/IRF9), as well as the antiviral, interferon-stimulated gene (ISG) MX Dynamin Like GTPase 1 (MX1)^33^. For MX1, we did not find a strong upregulation (Figure 6A). This is in line with the observation that the interferons (IFNs) α, β and γ included in our panel showed no upregulation. Also in bulk RNA-seq we did not find upregulation of interferons at any timepoint. For IRF9 and particularly IRF3 however, we observed clear indications of upregulation in keratinocytes early on, including cells not containing any viral RNA (Figure 6A). Interestingly, this effect was not detectable in bulk RNA-seq (Supplementary Figure 8B). As a confirmation, we found an upregulation of IRF3 also on the protein level, but only when keratinocytes were virus positive (Figure 6B). This suggests a delay of the IRF3 response on protein level, similar to our observation with IER3.

### HSV-1 induces a spatially coordinated expression of TNF family cytokines

In our bulk RNA-seq experiment we found that genes involved in response to TNF were overrepresented in the significantly upregulated genes at 4dpi (Figure 2B). The cytokines TNF (coding for TNFα) and TNFSF9 (coding for 4-1BBL) were significantly upregulated by HSV-1 infection in bulk- RNA-seq (Supplementary Figure 8B). Notably, spatial transcriptomics showed, that their expression is highly cell type-dependent and largely mutually exclusive, with TNF induced in keratinocytes (mainly stratum basale) and TNFSF9 in papillary fibroblasts with expression of both cytokines clearly depending on viral presence (Figure 7A). Their expression increased with increasing viral load up to a plateau level (Figure 7B). Published data from HSV-1 infected fibroblasts^23^ and HaCat keratinocytes^34^ support the findings of a cell type specific upregulation of TNFSF9 in fibroblasts and TNF in keratinocytes. Importantly, both the bulk RNA-seq data generated here in SkOs as well as the published bulk RNA-seq did not indicate that upregulation of TNF and TNFSF9 are results of HSV-1 induced read-in transcription from an upstream gene. To investigate, whether expression of TNF would lead to functional sensing in surrounding cells, we defined papillary fibroblasts proximal to TNF expressing cells (Supplementary Figure 8C) and analyzed them for induction of TNFα target genes (Figure 7D) based on a published RNA-seq data set generated in fibroblasts^35^. By calculating the differential expression in cells proximal vs. distal to the TNF-expressing cells we found that four of the 8 genes included in our panel were significantly upregulated especially in cells without viral gene expression (Figure 7D). By focusing on CXCL2, a TNFα-induced gene, we found a high number of cells expressing CXCL2 in the dermis close to TNF expressing keratinocytes (Figure 7C, upper row, last panel). To investigate whether this could also be caused by 4-1BBL (TNFSF9), we analyzed a region with high TNFSF9 but low TNF expression (Figure 7C, lower row). CXCL2 levels in these regions were lower, suggesting that the cell-type specific induction of the TNF gene leads to secretion of the TNFα cytokine and subsequently induction of target genes in the immediately surrounding cells.

**Figure 7:**
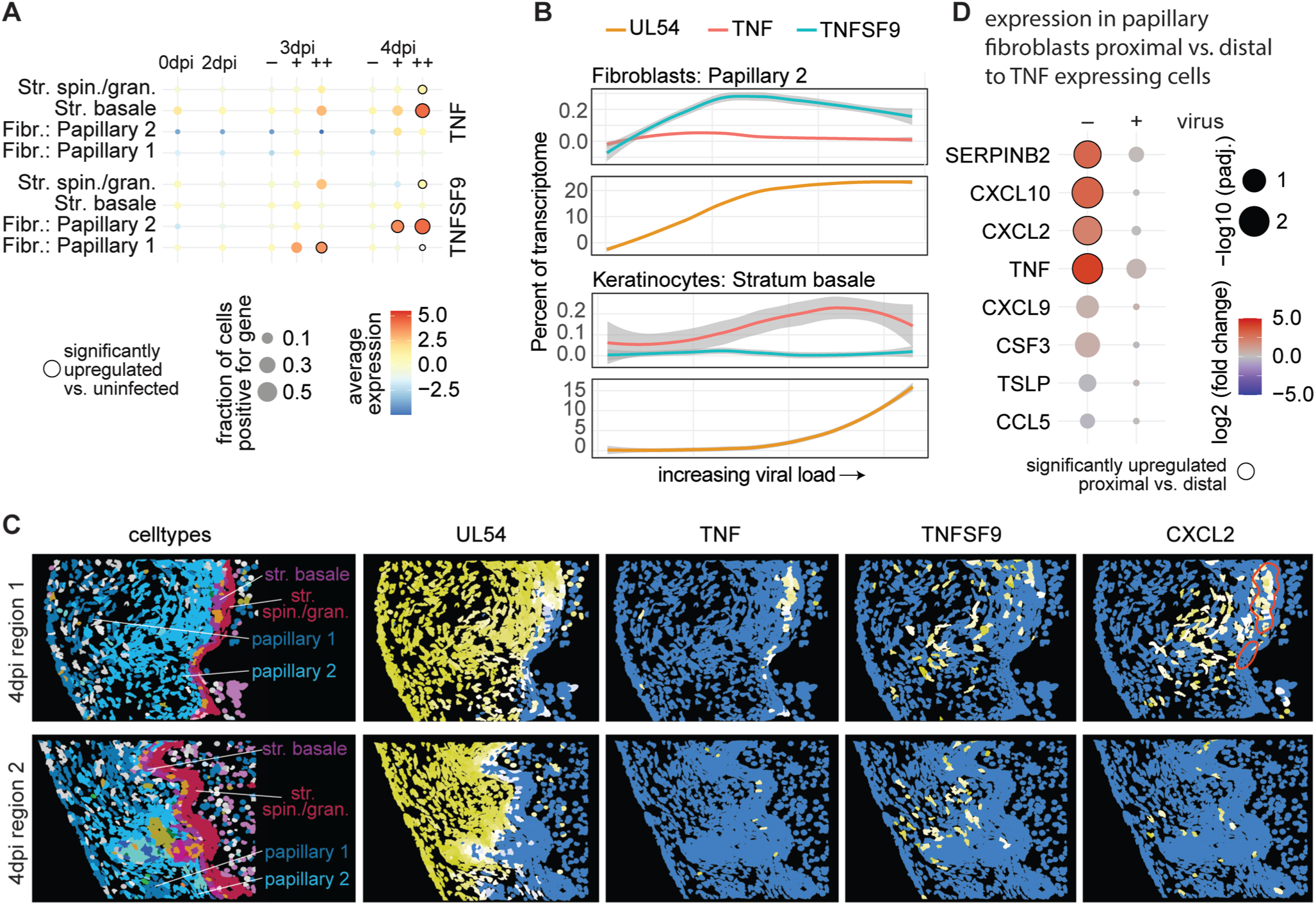
TNF induction leads to proximal expression of target genes. **A:** Expression values of the indicated genes in a subset of cell types. For uninfected and 2dpi SkOs, all cells were combined for the analysis. For 3dpi and 4dpi, cells are split up in those with no or only 1 viral count (-), and those with viral load below (+) and above (++) the median. The median viral load was calculated only within cells with more than 1 count. Dot sizes represent the fraction of cells having at least one count of the specified gene, the color the average expression. Black circles around the dots indicate significant differential expression in a pseudobulk comparison vs. all cells in uninfected cells (padj <0.05). **B:** Cells with at least 2 viral counts were ordered along viral load and binned with 20 cells per bin. Shown are the percentages of TNF, TNFSF9 and HSV-1 UL54 within the bin for papillary fibroblasts type 2 (top) and stratum basale keratinocytes (bottom). **C:** Shown are two regions of a 4dpi organoid, with cells colored by cell type or expression levels of the indicated genes. The cells encircled in red in the CXCL2 expression panel at 3dpi are expressing TNF. D: Differential expression of TNF target genes in papillary fibroblasts proximal vs. distal to TNF expressing cell groups.

## Discussion

In this study, we showed an infection of complex human hair bearing skin organoids with HSV-1. Due to the inside-out morphology of the organoids and the infection therefore starting from the dermis our model is closest to the reactivation scenario of HSV-1, when infectious particles are released from neuronal cells into the fibroblast layer of the dermis^36^. This makes our model highly relevant for investigating HSV-1 induced pathologies, since most of them are a result of reactivation and not primary infection. For instance, recurrent herpes labialis has an average incidence of about 1.6 per 1000 patients each year^37^.

When analyzing the susceptibility of the cells to HSV-1 in the SkO model we found that eventually at 8dpi all cell types in our model system were infected by HSV-1 except for the terminally differentiated keratinocytes, cornified structures in the HFs and chondrocytes in the tail. This is expected due to a lack of immune cells that could clear the infection. At the moment, we do not understand why chondrocytes are resistant to viral infection. A resistance of the stratum corneum against HSV-1 infection has been reported by many other studies using mouse models and *ex vivo* skin showing that the cornified keratinocyte layer prevents HSV-1 infection by the external surface^6,9^. The terminally differentiated keratinocytes of the epidermis in the skin organoid model express markers of the stratum corneum (LOR, KRT10), however, they do not eventually cornify since they are not exposed to air. Despite this difference their complete resistance to infection is well in line with findings from other models. Studies also showed that infection of all other layers of the epidermis by HSV-1 strongly depends on the infection route and the model system (dermal vs. epidermal, human vs. mouse, skin vs. mucosal skin)^38,39^. In human *ex vivo* skin wounding from the external side through different approaches leads to only limited infection of the epidermis^11,12^. Only infection from the dermal side by either removal of the dermis or deeper wounding through the epidermis into the dermis leads to efficient infection of the basal keratinocytes and spread to the higher differentiated keratinocyte layers^9,40^. In the SkO model the dermis was efficiently infected by HSV-1 and viral loads in papillary fibroblasts were much higher than in keratinocytes of the stratum basale and stratum spinosum/granulosum despite the presence of infectious cells in the neighborhood.

While the basal membrane as an extracellular matrix barrier likely slows down viral infection, we show evidence that the viral life cycle in keratinocytes is delayed compared to papillary fibroblasts. When we analyzed the spread of HSV-1 into HFs we observed that dermal sheath fibroblasts pose a barrier to HSV-1 infection of the more inner layers. The close physical proximity of dermal sheath fibroblasts to heavily infected papillary fibroblasts, in the absence of an intervening or expanded extracellular matrix, suggests that extracellular matrix structures are unlikely to account for the observed barrier function. Furthermore, the finding that dermal sheath cells are not infected less frequently but instead harbor a reduced viral load indicates that this phenotype is unlikely due to limited accessibility of a viral entry receptor, but rather reflects a delayed viral life cycle in this cell type. Our electron microscopy impressively shows that eventually almost all cell types in the HFs produce mature viral particles that were found intra- and extracellular. We hypothesize that the delayed HSV-1 infection in HFs could be important during reactivation when viral particles are released by the nerve endings of sensory neurons that innervate HFs. A restriction of the viral life cycle in dermal sheath fibroblasts could give immune cells time to counteract the infection. In the future, we would like to test this hypothesis by further analyzing the immune reaction in dermal fibroblasts and by including immune cells into the SkO model.

Also, we could show that the host response to HSV-1 infection in the SkO model is very cell type specific. We analyzed this on single cell level, but also in a spatial-temporal manner by analyzing cells at the infection front in the dermis. We found IER3 transcripts specifically upregulated in papillary fibroblasts at the leading edge of the infection. IER3 mRNA contains an AU-rich instability element (ARE) in its 3’UTR, a feature that promotes rapid mRNA degradation by deadenylation under normal conditions and stabilization by proinflammatory signaling like the p38 MAPK pathway^41,42^. Many studies have shown before in cell culture that the IER3 mRNA is stabilized by HSV-1^30,31^, in particular by ICP27^32^. Interestingly, transcript upregulation was transient, while the upregulated protein was very stable in the heavily infected cells in SkOs. A study using HeLa cells showed a transient induction of IER3 protein expression at 1 hour post infection (hpi) and a rapid decline already at 3hpi^31^. This could indicate a different turnover of the IER3 protein in tissue in comparison to a cell line.

The fact that we found a well known target of HSV-1 infection by our spatial temporal analysis shows the validity of the approach. It allows a comprehensive analysis of spatial transcriptomics data from infected tissues that goes beyond the discrimination between cell types and viral loads. However, IER3 was one of only three genes specifically upregulated at the infection front. This is likely an effect of the limited number of host genes included in our spatial transcriptomics experiment (470 genes) and we speculate that panels with more genes included will be even more informative to understand how a tissue reacts to a locally spreading infection. Another limitation of this approach is that for defining an infection front a certain number of cells from a specific cell type is needed, so we could perform this only for papillary fibroblasts of the dermis. Thus for lower abundant and spatially more distributed cells we compared uninfected cells and cells with low viral load and high viral load assuming that cells with a higher viral load have been infected for a longer time. We found that IRF3 is upregulated at transcript and protein level in keratinocytes of the stratum basale. Similar to IER3 we find a delayed effect of protein induction in comparison to the transcripts and no induction of IRF3 on protein level in virus negative cells. Since we do not find induction of IFNα, β or γ, or the ISGs included in our panel we can only speculate which downstream signaling might be induced by IRF3. An indication for a potential antiviral effect is that IRF3 protein is mainly upregulated in infected keratinocytes of the stratum basale where we also found lower viral loads in comparison to fibroblasts of the dermis.

The host response to HSV-1 infection in SkO is further dominated by inflammation. Spatial transcriptomics showed that TNF (coding for TNFα) was induced in keratinocytes of the stratum basale, and TNFSF9 (coding for 4-1BBL) in papillary fibroblasts. We also observed induction of TNF target genes in surrounding cells and a potential priming of adjacent uninfected cells, indicating that gene activation is indeed functional. Induction of TNF and subsequent tissue damage was previously associated with HSV-1 in cerebral infections^43^. In *ex vivo* mouse whole corneal culture HSV-1 spread was reduced by TNF treatment but also resulted in increased cell death^44^. In skin it is likely that the strong inflammatory reaction induced by the different TNF cytokines helps to limit viral replication in the case of reactivation in a cell type specific manner. The drawback is the induced tissue damage, but it also offers opportunities to specifically reduce HSV-1 induced inflammation and tissue damage by treatment with inhibitors for TNF or TNFSF9.

In summary, we show here that SkOs are a physiologically highly relevant model system for HSV-1 infection in the skin. The combination with single-cell-resolution spatial transcriptomics enabled an unprecedented detailed analysis of viral gene expression and host response. Compared with *ex vivo* skin models, SkOs offer key advantages, including the feasibility of genetic manipulation and prolonged maintenance in culture. These features make SkOs particularly well suited for investigating HSV mutants with growth defects, for studies of viral persistence and latency as well as drug testing. The limitations of the SkO model system lie in the lack of immune cells and vascularization, but a recent study reported the successful incorporation macrophages and improved angiogenesis in SkOs^21^. Building on these advances, future work will aim to further refine this model for HSV-1 and deepen our understanding of HSV-1 pathogenesis in the skin.

## Material and Methods

### Skin organoid generation

Skin organoids (SkOs) were generated from the human iPS cell line UKEi001-A (RRID:CVCL_A8PR). All procedures involving hiPSC lines were approved by the local ethics committee in Hamburg (Az PV4798, 28.10.2014). HiPSCs were cultivated in six-well plates coated with Geltrex (Thermo Fisher Scientific, cat. no. A1413302) in StemFlex medium (Gibco, cat. no. A3349401) under standard conditions (37°C, 5% CO₂). Generation of SkOs was performed according to the protocol by Lee and colleagues^19^ with hiPSCs between passages 31-39. Only SkOs showing correct differentiation as determined by the appearance of HFs (HF) at 110 to 120 days post differentiation were used for further experiments.

### Virus infection

HSV-1 17 CMV-IEproEGFP expressing EGFP under the control of the CMV IE promoter was described before^45^. The virus was propagated on Vero cells and the infection titer determined by plaque assay. Medium on SkOs was exchanged and infection was performed by adding 800 PFU per SkO to the medium. After 12hrs the medium was exchanged with standard SkO growth medium and afterwards complete medium exchange was performed every day.

### Immunostaining and microscopy

For immunohistochemistry SkOs were fixed in 4% phosphate buffered formalin over night at 4°C and then transferred to PBS. Afterwards they were protected in 3% agarose, dehydrated using a Leica ASP300S tissue processor, and embedded in paraffin. Organoid paraffin sections were cut at 3 μm and stained with hematoxylin and eosin (H&E) according to standard procedures. For immunohistochemical staining the sections were processed as following: After dewaxing and inactivation of endogenous peroxidases (3% hydrogen peroxide), antibody specific antigen retrieval was performed using the Ventana BenchMark XT autostainer (Ventana, Tuscon, Arizona, USA). Sections were blocked and afterwards incubated with the primary antibodies, see Table S1. For detection of specific binding and DAB staining, the UltraView Universal DAB Detection Kit (Roche, #760-500) was used, which contains both, anti-mouse and anti-rabbit secondary antibody. Counter staining and bluing were performed with Hematoxylin (Ventana Roche, # 760-2021) and Bluing Reagent (Ventana Roche, # 760-2037) for 4 min. Subsequently, stained sections were mounted in mounting medium.

Pictures of IHC staining were acquired with the Sysmex Pannoramic MIDI II slide scanner. Image cropping was performed with QuPath0.4.3 and pictures assembled with Adobe Illustrator.

For whole mount staining of SkOs the qCe3D method was used according to Lee and colleagues^19^ with the following adjustments: Clearing was performed for 48hrs and stained and cleared SkOs were mounted in iSpacer (SUNJinLab) 4 well imaging spacers. For antibodies see Supplementary Table S1. Imaging was performed with a Leica TCS SP8 X confocal microscope. Images were processed with Imaris and ImageJ 2.14 and pictures assembled with Adobe Illustrator.

### Electron microscopy

SkOs were fixed overnight in 2% paraformaldehyde and 2.5% glutaraldehyde, washed in PBS, and post-fixed in 1% osmium tetroxide for 60 min. Following additional rinses in PBS and double-distilled water, samples were stained with 1% aqueous uranyl acetate for 60 min, dehydrated through a graded ethanol series, and embedded in Epon (Carl Roth). Ultrathin sections (50 nm) were prepared using an Ultracut microtome (Leica Microsystems) and post-stained with 1% uranyl acetate in ethanol. Transmission electron micrographs were acquired on a FEI Tecnai G20 Twin operated at 80 kV and equipped with a 2K CCD Veleta camera (Olympus SIS). For correlating TEM data across entire organoids, serial sections were mounted on silicon wafers and imaged using a Tescan Clara SEM with backscattered-electron detection at 2.8 kV.

### Bulk RNA-seq

For bulk RNA-sequencing 4 SkOs per condition (uninfected, 2 days post infection (dpi), 4dpi, 6dpi, 8dpi) were used. Total RNA was isolated individually from each SkO by adding QIAzol lysis reagent (Qiagen) and disrupting the organoid by grinding with a pestle. The lysate was homogenized with a QIAshredder (Qiagen). After one round of chloroform extraction, RNA was cleaned up with the RNeasy MinElute Cleanup Kit (Qiagen). DNA was two times digested with the DNA-free DNA removal kit (Thermo Fisher Scientific). RNA concentration was determined with the Qubit RNA broad range assay (Thermo Fisher Scientific) and equal amounts of RNA from two SkOs of the same condition were pooled resulting in duplicates for each condition for sequencing. RNA integrity was determined via a Bioanalyzer RNA 6000 Pico Assay (Agilent, 5067-1514). RIN values were in the range between 5.5 and 9.2. From each pooled sample 520 ng total RNA was subjected to rRNA depletion (following the Lexogen RiboCop rRNA Depletion Kit for Human/Mouse/Rat V2 protocol, catalog number: 144.24) and further processed via the RNA-Seq V2 Library Prep Kit with UDIs (Lexogen, catalog number: 175.96) in the short insert size variant (RTM) according to manufacturer’s instructions. Libraries were quality controlled on a Bioanalyzer High Sensitivity DNA Assay (Agilent, 5067-4626) and were sequenced on an Illumina NextSeq 500 instrument with NextSeq 500/550 High Output Kit v2.5 (75 Cycles, product number: 20024906) in a single-read mode (1 x 83 bp). After demultiplexing via bcl2fastq, 20.7 – 28.6 million (M) read pairs assigned to each sample. All samples passed quality control by fastqc and were subjected to downstream analysis. Processing was done with the nf-core/RNA-seq pipeline (v3.12.0) within the nf-core framework^46^. The pipeline was executed with Nextflow v23.04.1^47^, utilizing STAR (v2.6.1d)^48^ for alignment to the GRCh38 human genome and Human herpesvirus 1 genome (X14112.1), as well as Salmon (v1.10.1)^49^ for transcript-level quantification based on STAR’s alignment output. To eliminate duplicate reads, the --with_umi option was enabled. Gene-level quantification was derived from transcript-level estimates using the R package tximport (v1.34.0)^50^, which sums transcript abundances for each gene. The generated raw gene-level counts were normalized and variance-stabilized using the regularized log transformation (rlog) from DESeq2 (v1.34.0)^51^, addressing the variance-over-mean trend. Principal component analysis (PCA) confirmed high reproducibility across biological replicates. Differential expression analysis was performed using DESeq2, identifying genes as significantly differentially expressed (DEGs) if they exhibited a fold change ≥ 2 and an adjusted P-value ≤ 0.05. The normalized rlog-transformed counts were then used for clustering and visualization. An overrepresentation analysis of differentially expressed host genes (padj. ≤0.05 and log2FC ≥1/≤-1) was performed with WebGestalt^52^. GO terms associated with “Biological processes noRedundant” were selected with an FDR ≤0.01.

### Spatial transcriptomics experiments

SkOs were imbedded in paraffin as described for IHC staining. 3 SkOs for each condition were assembled in one agarose mold before embedding all conditions (uninfected/2dpi/3dpi/4dpi for the main experiment, uninfected/2dpi/6dpi for the pilot experiment). *In situ* RNA measurements were performed using the Xenium system (10x Genomics). The pilot experiment was done at the MDC/BIH genomics platform using nuclear expansion based on DAPI staining for cell segmentation. The main experiment was performed at the 10xGenomics facility in Stockholm using multimodal segmentation (nuclear stain, RNA and protein-based cell staining, cell surface staining). For both experiments, the human multi tissue gene panel was used, supplemented by the 100 genes custom panel CZBXK3 (Supplementary Table 1). Sections of 5μm thickness were placed on a Xenium slide according to the manufacturer’s protocol, with drying at 42°C for 3 hours and overnight placement in a desiccator at room temperature, followed by deparaffinization and permeabilization to make the mRNA accessible. The Probe Hybridization Mix was prepared according to the user guide (CG000582, Rev D, 10x Genomics). The staining for Xenium was performed using Xenium Nuclei Staining Buffer (10x Genomics product number: 2000762) as a part of the Xenium Slides & Sample Prep Reagents Kit (PN-1000460). Following the Xenium run, Hematoxylin and Eosin (H&E) staining was performed on the same section according to the Post-Xenium Analyzer H&E Staining user guide (CG000613, Rev B, 10x Genomics).

### Spatial transcriptomics analysis

The spatial transcriptomics data was analyzed in R using the VoltRon package^53^, as well as packages from tidyverse^54^. Other packages used include DESeq2^51^, apeglm^55^, sf^56^ and concaveman^57^. Briefly, we first assessed cell type identity only on cellular transcriptomes, i.e. without taking spatial information into account (Supplementary Table 2 and see Supplementary Note for more information). Viral load was defined as percent viral counts (i.e., sum of the five viral genes UL54, US1, UL27, UL29, LAT) of all counts per cells. Cells with 10 or less cellular RNA counts were filtered out. Organoid boundaries and distances from these boundaries were identified using the concaveman/st_boundary/st_distance function from the concaveman^57^ and sf^56^ R packages. To classify cells based on the number of infectious cells in the neighborhood, the absolute number of cells with more than 20% viral load and within 50µm (centroid to centroid distance) was counted for every cell. Differential expression using DESeq2^51^ was calculated as pseudo-bulk per cell type, i.e., counts were summed up per group (uninfected and 2dpi all cells, 3 and 4dpi either cells with 1 or less viral count “-”, viral load below “+”, or above the median “++” within the group). The infection front analysis was initiated by clustering papillary fibroblasts with at least 2 viral counts using only genes that are differentially expressed upon infection. Cells of cluster 1, with the lowest viral load, which were in groups of at least 5 cells within 150 micrometers, were defined as the infection front. Cells with less than 2 viral counts and within 50µm of these cells were classified as “uninfected next to infection front”. For the TNF proximity analysis, cells were defined as “TNF expressing”, if at least 3 cells with at least 1 TNF gene count were not more than 30 µm apart from each other, and the proximal area defined at 280 µm around these cells. ChatGPT (OpenAI, https://chat.openai.com, model 5) assisted in writing R code.

## Data and code availability

Bulk RNA-seq data were deposited at NCBI under BioProject accession number PRJNA1378132, and the raw sequencing reads are available in the NCBI Sequence Read Archive (SRA) under accession numbers SRR36379728–SRR36379737. Spatial transcriptomics data are available through NCBI GEO (accession number GSE313919). Code for spatial transcriptomics and bulk RNA-seq data analysis is available at https://github.com/LandthalerLab/SkinOrganoids_HerpesSimplexVirus1.

## Acknowledgements

The authors thank Robert Shelansky, Aishwarya Konnur, Jana Lalakova, Morgane Rouault (10xGenomics) for performing the main spatial experiment in this manuscript and for technical support. Vera Wolf, Kathrein Permien (Charité) and Kristin Hartmann (Core Facility Experimental Histo-Pathology/UKE) provided technical assistance. We thank the UKE imaging facility (umif) for technical support in microscopy. We thank Claudia Schmidt, Lisann Röpke, Josephine Bonath and Oula Moshilam (UKE) for technical assistance in iPSCs and organoid culture. We thank Patrick Blümke (Technology Platform Next Generation Sequencing, LIV) for technical support in bulk RNA-seq.

This work was supported by the Deutsche Forschungsgemeinschaft (DFG; German Research Foundation) in the framework of the Research Unit FOR5200 DEEP-DV (443644894) project, CZ 320/1-1, GR 3318/5-1, LA 2941/18-1, FR 2938/11-1 and FI 782/7-1 to M.C.-S., A.G., M.L., C.C.F. and N.F., respectively. M.C.-S. was supported by PIER (Partnership of Universität Hamburg and DESY) project PIF-2023-13.

## Contributions

Conceptualization: E.W., S.A., A.G., M.L., N.F., M.C.-S

Methodology:. E.W., S.A., A.W.,M.C.-S.

Investigation: E.W.,S.A., A.W., N.B., S.K., R.R., C.S., H.R., M.C.-S.

Data analysis: E.W., C.C.F., J.H., M.C.-S.

Resources (reagents and techniques): E.W., A.M., I.P., J.A., T.C., W.H., A.H., A.G., M.L., N.F, M.C.-S.

Writing – original draft: E.W., M.C.-S.

Funding acquisition: A.G., C.C.F., M.L., N.F., M.C.-S.

All authors discussed the results, contributed to manuscript editing, and approved the final version.

## Competing interests

The authors declare no competing interests.

**Supplementary Figure 1: A:** Brightfield images of SkOs from day −2 (d-2) of differentiation to 126 days (d126). White scale bars represent 200µm, black scale bars 500µm. **B:** Maximum intensity projections from Z-stacks of overview images from WMI staining of SkOs at 120 days of differentiation. Scale bars 500µm. **C:** Zoom-ins of WMI staining. Scale bars 50µm. **D:** IHC staining of SkOs at 120 days of differentiation. Boxes in overview images indicate areas of higher magnification shown at the right. Scale bars represent 500µm in overview images and 50µm in zoom-ins. H&E: hematoxylin and eosin stain. Hoechst: nuclear stain. Ki67: proliferating cells. KRT15: basal layer of epidermis, outer root sheath, medulla. KRT17: epidermis, outer root sheath of HFs. KRT20: Merkel cells. KRT71: inner root sheath of HFs. LHX2: bulge region, hair germs, hair pegs, hair placodes. NEFH: sensory neurons, large soma neurons. PDGFRα: dermal fibroblasts, dermal papilla. S100β: Schwann cells, satellite glial cells, adipocytes, melanocytes. SCD1: adipocytes, sebaceous glands. SOX2: dermal condensates, dermal papilla, melanocytes, Merkel cells. TUBB3: neuronal cells, Schwann cells.

---

**Supplementary Figure 2: A:** IHC staining against GFP of 120 days old SkOs infected with HSV-1 expressing GFP from CMV-IE promoter. Boxes in overview images indicate areas of higher magnification shown below. Scale bars 500µm in overview images and 20µm in zoom-ins. Upper picture of zoom-ins shows epidermis on the top and dermis below. Lower picture in zoom-ins shows hair follicle. **B:** Heatmap of bulk RNA-seq of HSV-1 infected SkOs showing deregulated genes of HSV-1 infected SkOs at 2, 4, 6, and 8dpi vs uninfected SkOs in the individual replicates. **C:** Volcano plots of bulk RNA-seq showing all differentially expressed host genes at 2, 4, 6, and 8dpi vs uninfected SkOs.

**Supplementary Figure 3:** Spatial transcriptomics of HSV-1 infected SkOs. Per timepoint (uninfected, 2dpi, 3dpi, 4dpi) 3 SkOs were measured. **A:** Overview of all SkOs analyzed by spatial transcriptomics with cells colored by cell types. **B:** Cells as in A, colored by normalized expression of HSV-1 UL54 RNA. **C:** Number of cells per organoid replicate. **D:** Violin plot depicting distribution of detected genes and RNA counts for the major cell types.

**Supplementary Figure 4. A:** Papillary fibroblasts type 1 cells were binned according to their distance from the organoid boundary (µm) in 10 bins of equal size. Other cell types as indicated were sorted into the bins with the same distance range as the respective papillary fibroblasts type 1 bin. By that, the distance distribution per bin is the same for all cell types (top panel), but the fraction of cells per bin is equal only for papillary fibroblasts type 1 in all bins (bottom panel), since the other cell types in general reside further inside the organoid. **B:** Top panel, fraction of cells with at least two viral counts per distance bin. Bottom panel, viral load (percent viral transcripts) in the different cell types per distance bins.

**Supplementary Figure 5: A:** Cells from SkOs at 4dpi were binned by cell type and by the number of infectious cells (defined as >20% viral transcripts) in their neighborhood (within 50µm). Shown is the viral load (percent viral transcripts) per group as boxplots. The middle line in the boxplot displays the median, the box indicates the first and third quartile, whiskers the 1.5 interquartile range (IQR). Outliers beyond are marked by single dots. **B:** Shown is one replicate of a HSV-1 infected SkO at 4dpi (bottom right of the 4dpi organoids in Supplementary Figure 3A). Dermal sheath cells within 50µm of infectious cells (defined as >20% viral transcripts) are labelled in dark blue. Cells of hair follicle cell types (dermal papilla, bulge region, IRS, ORS, matrix, melanocytes) within 50µm of these dermal sheath cells are labelled in medium pink. Other cell types (i.e., non-hair follicle cell types, thus outside of HFs, such as papillary fibroblasts) are labelled in petrol. **C:** DAPI, cell surface protein and interior protein staining on the Xenium slide. Surrounded with orange blue are dermal sheath cells as in Figure 4D.

**Supplementary Figure 6:** Scanning electron microscopy (SEM) of HFs from HSV-1 infected SkOs at 3dpi and 4 dpi. **A:** Overviews showing the scanned areas. Black boxes indicate areas scanned with higher resolution. White boxes with numbers indicate areas of zoom-ins shown in B. pFib: papillary fibroblasts; DS: dermal sheath; GM: glassy membrane; ORS: outer root sheath; parts of inner root sheath (IRS): Cp: companion layer, He: Henle‘s layer, Hu: Huxley‘s layer, Ci: IRS cuticle; Mtx: matrix; DP: dermal papilla. Scale bar: 50µm. **B:** Zoom-ins on areas indicated in part A. Red arrows indicate viral particles. Yellow boxes indicate areas of zoom-ins showing extracellular viral particles and a nuclear replication compartment with A, B and C capsids. RC: viral replication compartment; Ex: extracellular space, N: nucleus, C: cytoplasm. Scale bar: 1µm; 200nm in zoom-ins.

**Supplementary Figure 7: A:** Two-dimensional projection of papillary fibroblasts as in Figure 5A, with cells colored by expression levels of the indicated host genome transcripts. **B:** Papillary fibroblasts were categorized in “infection front” (cluster 1 in Fig. 5AB), “uninfected next to infection front”, “infected next to infection front”, “later in infection” and “no virus” (i.e., without viral RNA and further away from the infection front). Top: two-dimensional projection as in A, bottom: the same region as in Figure 5B, both with cells colored by these categories. **C:** Differential gene expression in papillary fibroblasts in the cell categories shown in B at 3dpi vs uninfected. Shown are the genes with padj < 0.05 in at least one category. **D:** Fold changes in the bulk RNA-seq data at the indicated timepoints vs. uninfected organoids for IER3 and selected induced host genes. Significantly upregulated genes have padj <0.05.

**Supplementary Figure 8: A:** Expression values of the indicated genes in a subset of cell types. For uninfected and 2dpi SkOs, all cells were combined for the analysis. For 3dpi and 4dpi, cells are split up in those with no or only 1 viral count (-), and those with viral load below (+) and above (++) the median. The median viral load was calculated only within cells with more than 1 count. Dot sizes represent the fraction of cells having at least one count of the specified gene, the color the average expression. Black circles around the dots indicate significant differential expression in a pseudobulk comparison vs. all cells in uninfected cells (padj <0.05). **B:** Fold changes in the bulk RNA-seq data at the indicated timepoints vs. uninfected organoids for IRF3, TNF and TNFSF9. Significantly upregulated genes have padj <0.05. **C:** Shown is one replicate of a HSV-1 infected SkO at 4dpi (top left of the 4dpi organoids in Supplementary Figure 3A and Figure 7C). Cells are colored by TNF expressing cells, proximal and distal cells as indicated.

