## Supplementary Figure 1 for "Complex Human hear bearing Skin Organoids reveal Cell Type Specific Susceptibility and Innate Immune Responses to Herpes Simplex Virus 1"

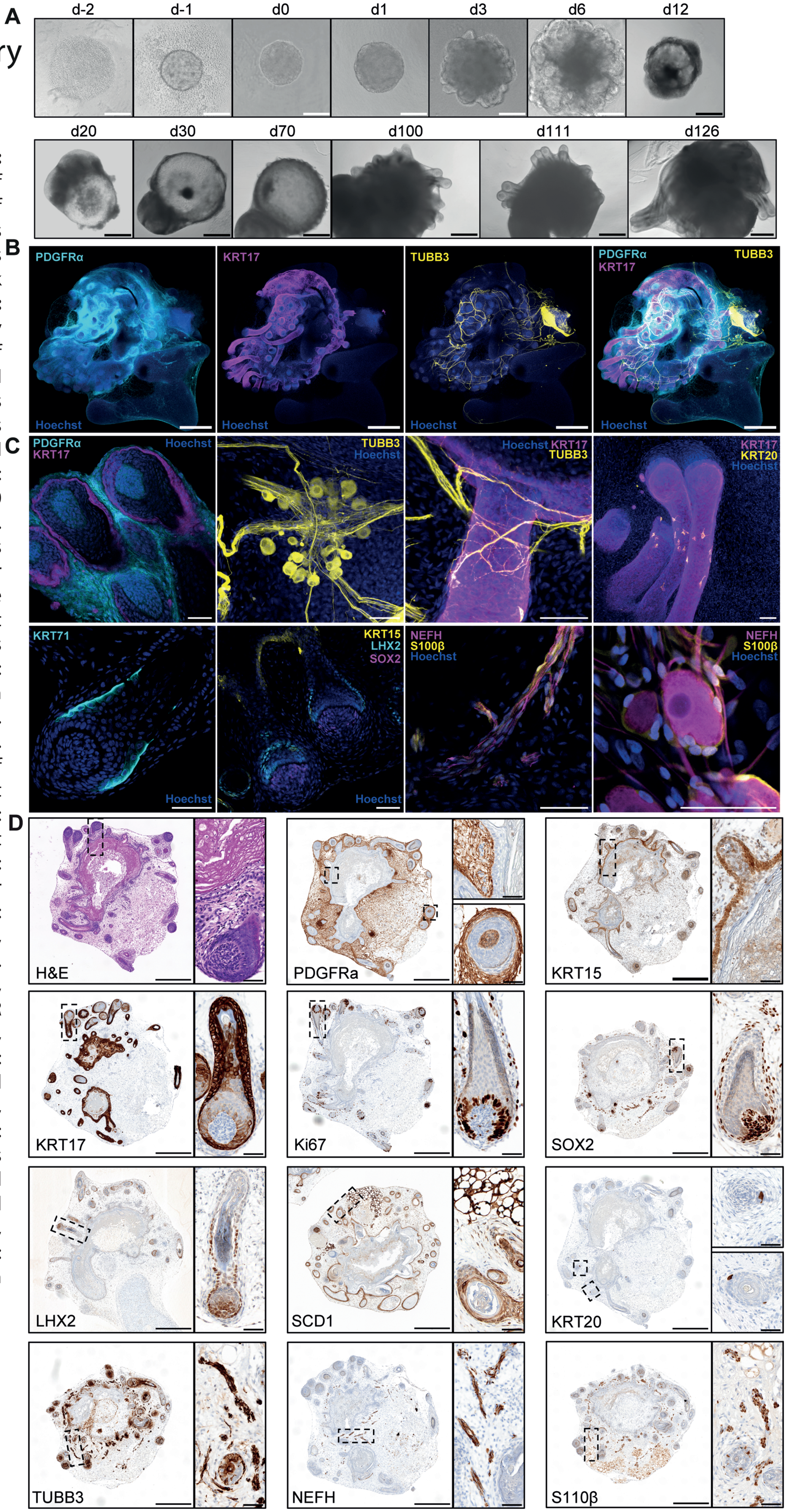


**Supplementary Figure 1:** **A:** Brightfield images of SkOs from day -2 (d-2) of differentiation to 126 days (d126). White scale bars represent 200µm, black scale bars 500µm. **B:** Maximum intensity projections from Z-stacks of overview images from WMI staining of SkOs at 120 days of differentiation. Scale bars 500µm. **C:** Zoom-ins of WMI staining. Scale bars 50µm. D: IHC staining of SkOs at 120 days of differentiation. Boxes in overview images indicate areas of higher magnification shown at the right. Scale bars represent 500µm in overview images and 50µm in zoom-ins. H&E: hematoxylin and eosin stain. Hoechst: nuclear stain. Ki67: proliferating cells. KRT15: basal layer of epidermis, outer root sheath, medulla. KRT17: epidermis, outer root sheath of HFs. KRT20: Merkel cells. KRT71: inner root sheath of HFs. LHX2: bulge region, hair germs, hair pegs, hair placodes. NEFH: sensory neurons, large soma neurons. PDGFR α: dermal fibroblasts, dermal papilla. S100β: Schwann cells, satellite glial cells, adipocytes, melanocytes. SCD1: adipocytes, sebaceous glands. SOX2: dermal condensates, dermal papilla, melanocytes, Merkel cells. TUBB3: neuronal cells, Schwann cells.
