## Supplementary Figure 2 for "Complex Human hear bearing Skin Organoids reveal Cell Type Specific Susceptibility and Innate Immune Responses to Herpes Simplex Virus 1"

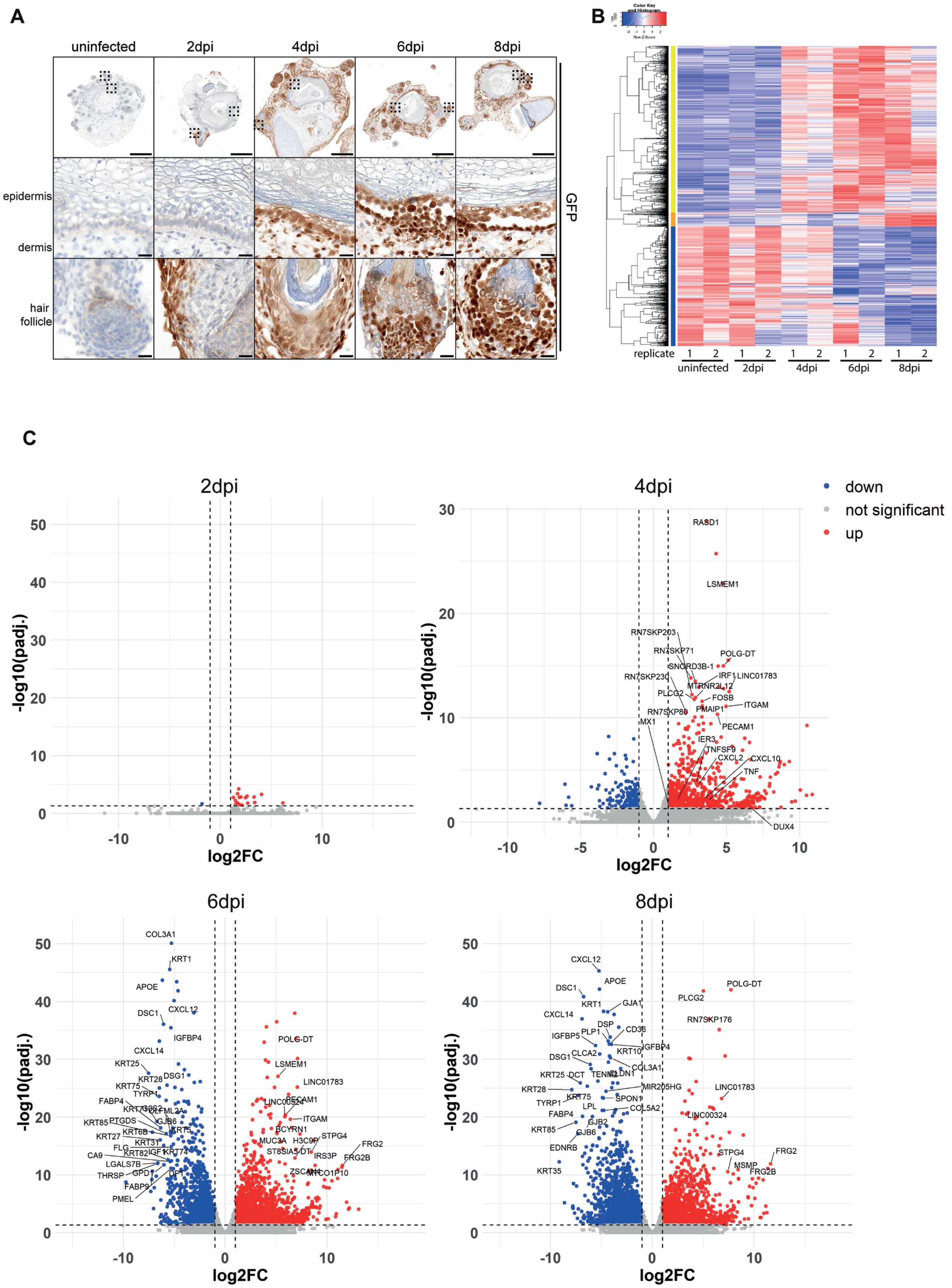


**Supplementary Figure 2: A:** IHC staining against GFP of 120 days old SkOs infected with HSV-1 expressing GFP from CMV-IE promoter. Boxes in overview images indicate areas of higher magnification shown below. Scale bars 500µm in overview images and 20µm in zoom-ins. Upper picture of zoom-ins shows epidermis on the top and dermis below. Lower picture in zoom-ins shows hair follicle. **B:** Heatmap of bulk RNA-seq of HSV-1 infected SkOs showing deregulated genes of HSV-1 infected SkOs at 2, 4, 6, and 8dpi vs uninfected SkOs in the individual replicates. **C:** Volcano plots of bulk RNA-seq showing all differentially expressed host genes at 2, 4, 6, and 8dpi vs uninfected SkOs.
