## Supplementary Figure 3 for "Complex Human hear bearing Skin Organoids reveal Cell Type Specific Susceptibility and Innate Immune Responses to Herpes Simplex Virus 1"

### Supplementary Figure 3, part 1

A

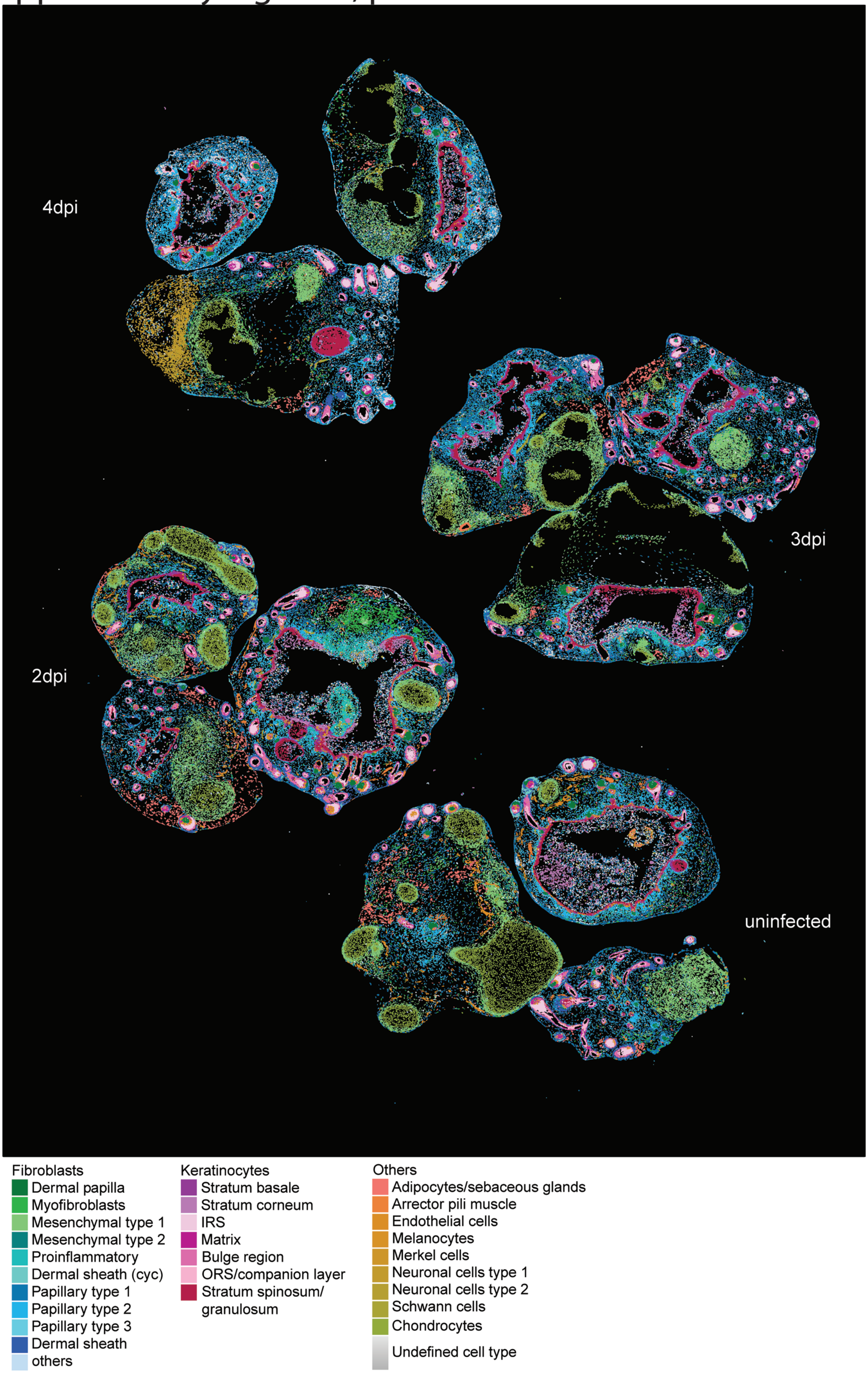


**Supplementary Figure 3: Spatial transcriptomics of HSV-1 infected SkOs.** Per timepoint (uninfected, 2dpi, 3dpi, 4dpi) 3 SkOs were measured. **A:** Overview of all SkOs analyzed by spatial transcriptomics with cells colored by cell types. **B:** Cells as in A, colored by normalized expression of HSV-1 UL54 RNA. **C:** Number of cells per organoid replicate. **D:** Violin plot depicting distribution of detected genes and RNA counts for the major cell types.

Supplementary Figure 3, part 2

**B**

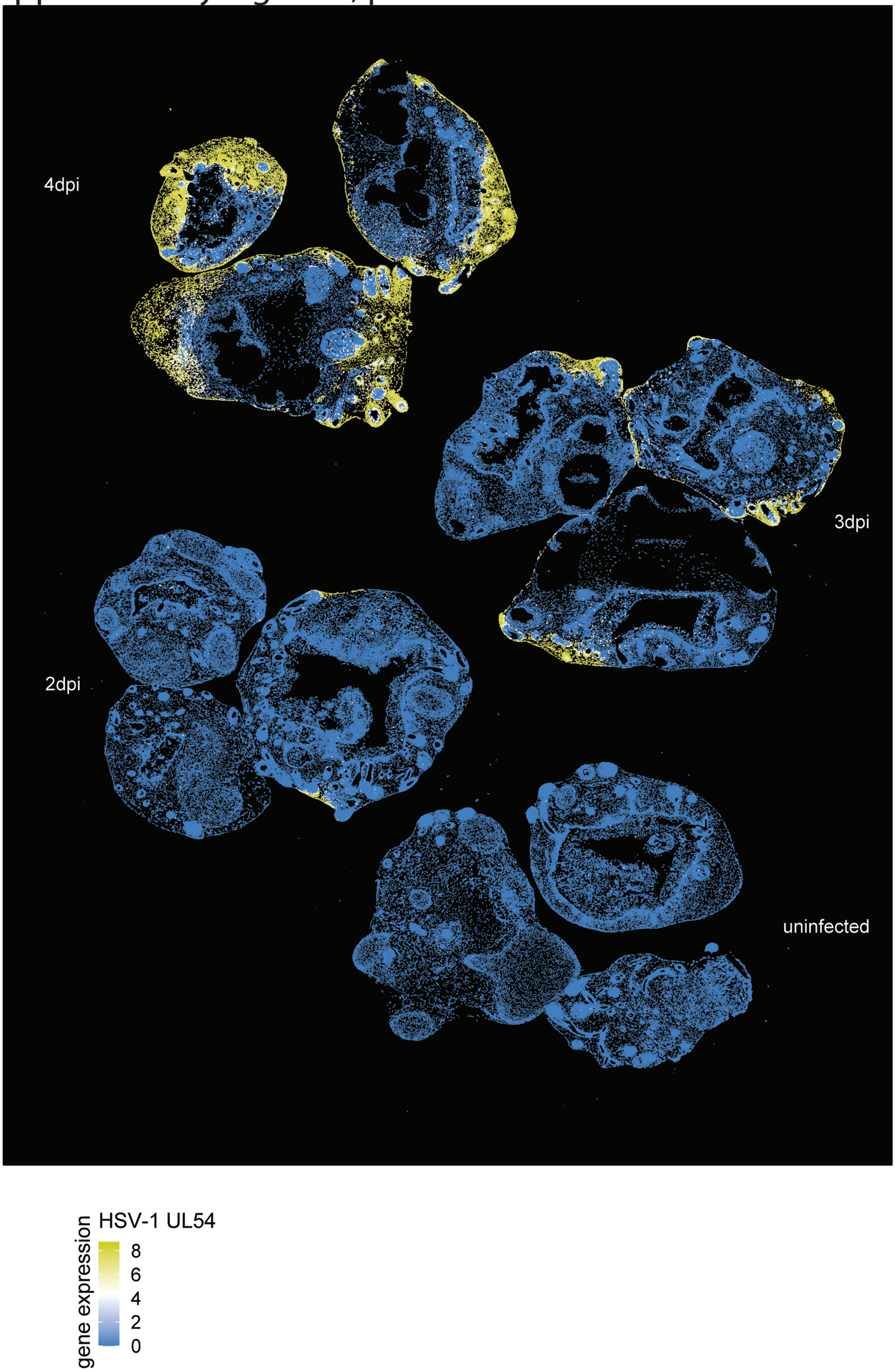

### Supplementary Figure 3, part 3

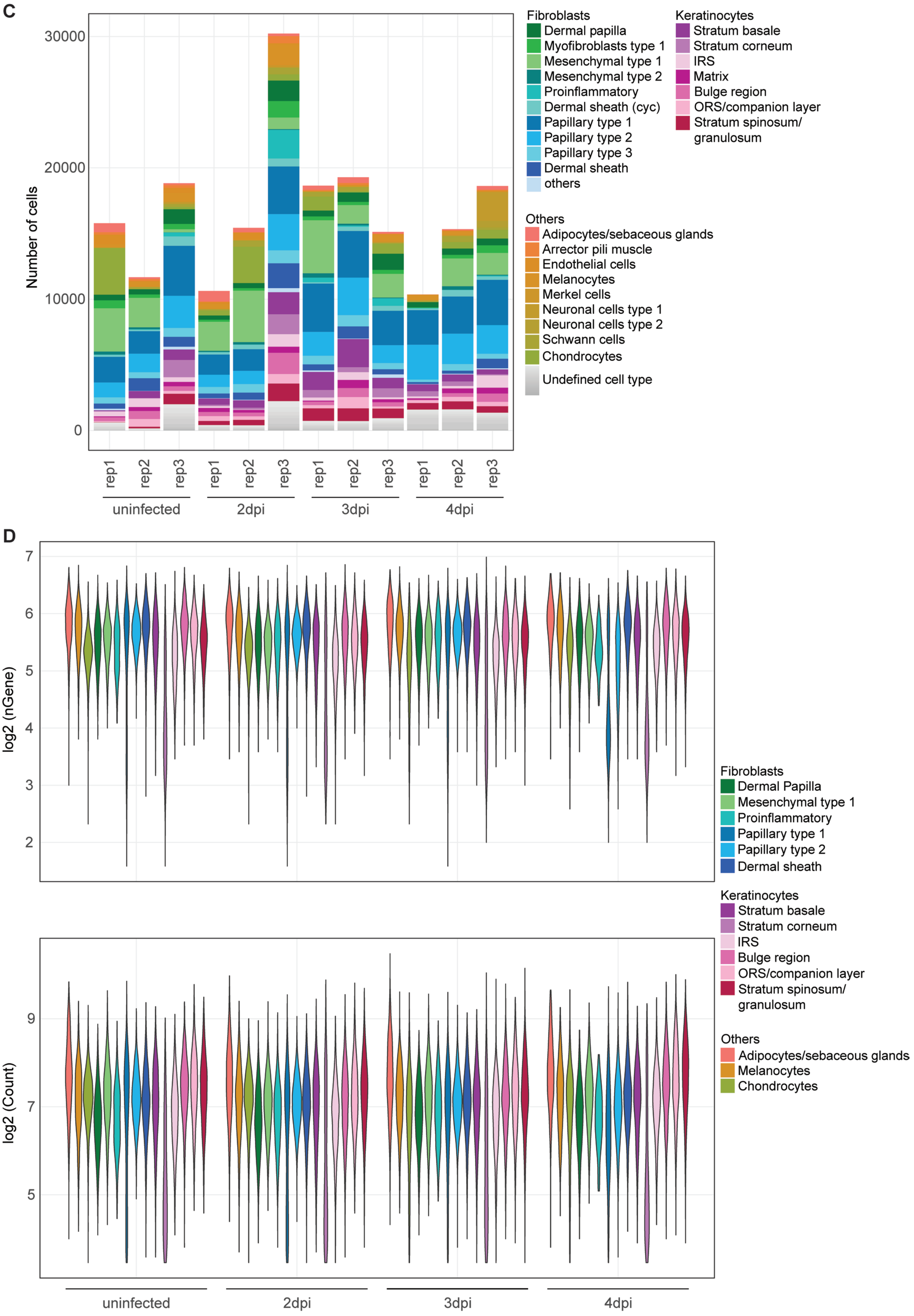
