## Supplementary Figure 4 for "Complex Human hear bearing Skin Organoids reveal Cell Type Specific Susceptibility and Innate Immune Responses to Herpes Simplex Virus 1"

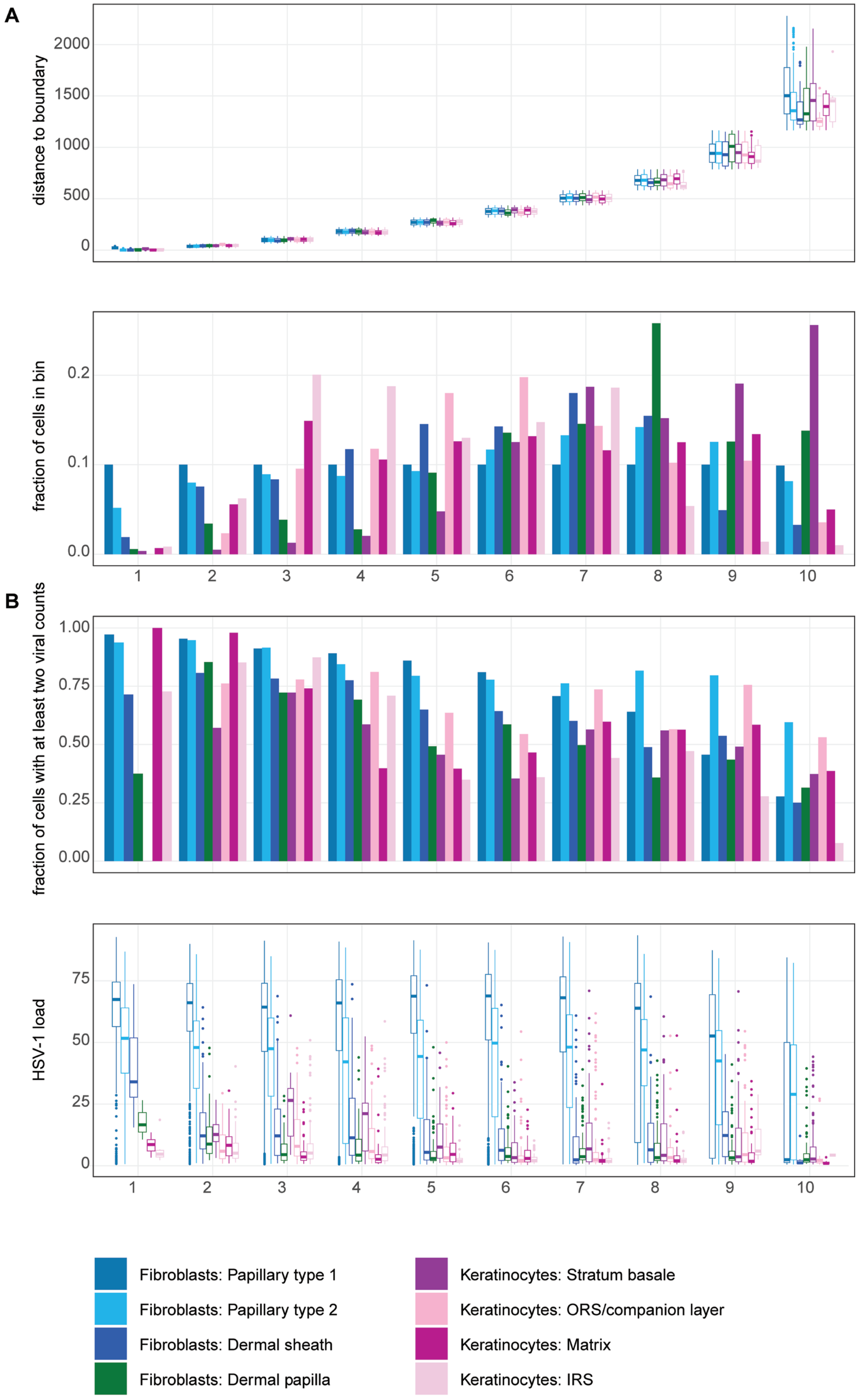


**Supplementary Figure 4. A:** Papillary fibroblasts type 1 cells were binned according to their distance from the organoid boundary (µm) in 10 bins of equal size. Other cell types as indicated were sorted into the bins with the same distance range as the respective papillary fibroblasts type 1 bin. By that, the distance distribution per bin is the same for all cell types (top panel), but the fraction of cells per bin is equal only for papillary fibroblasts type 1 in all bins (bottom panel), since the other cell types in general reside further inside the organoid. **B:** Top panel, fraction of cells with at least two viral counts per distance bin. Bottom panel, viral load (percent viral transcripts) in the different cell types per distance bins.
