## Supplementary Figure 5 for "Complex Human hear bearing Skin Organoids reveal Cell Type Specific Susceptibility and Innate Immune Responses to Herpes Simplex Virus 1"

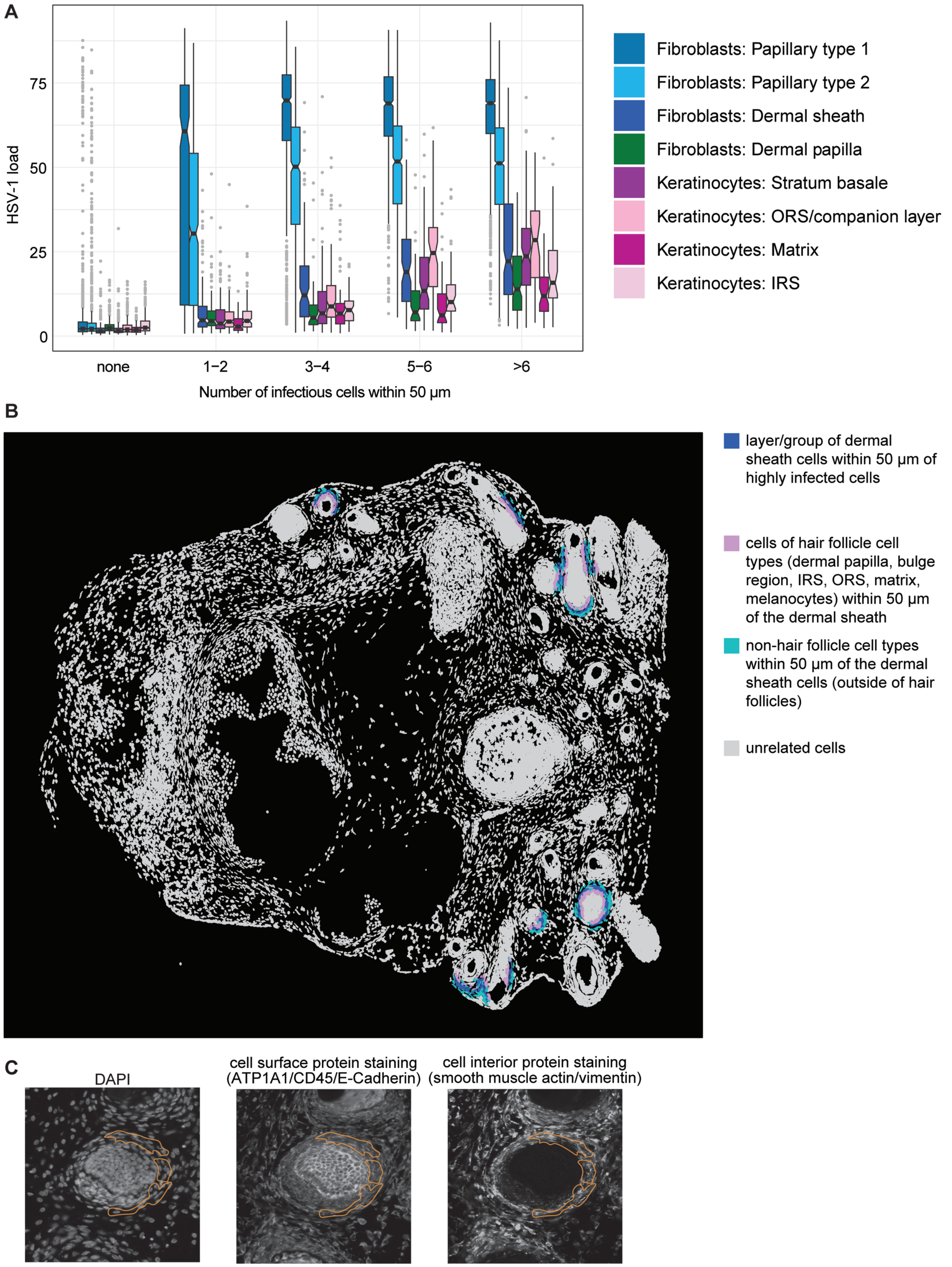


**Supplementary Figure 5: A:** Cells from SkOs at 4dpi were binned by cell type and by the number of infectious cells (defined as >20% viral transcripts) in their neighborhood (within 50µm). Shown is the viral load (percent viral transcripts) per group as boxplots. The middle line in the boxplot displays the median, the box indicates the first and third quartile, whiskers the 1.5 interquartile range (IQR). Outliers beyond are marked by single dots. **B:** Shown is one replicate of a HSV-1 infected SkO at 4dpi (bottom right of the 4dpi organoids in Supplementary Figure 3A). Dermal sheath cells within 50µm of infectious cells (defined as >20% viral transcripts) are labelled in dark blue. Cells of hair follicle cell types (dermal papilla, bulge region, IRS, ORS, matrix, melanocytes) within 50µm of these dermal sheath cells are labelled in medium pink. Other cell types (i.e., non-hair follicle cell types, thus outside of HFs, such as papillary fibroblasts) are labelled in petrol. **C:** DAPI, cell surface protein and interior protein staining on the Xenium slide. Surrounded with orange blue are dermal sheath cells as in Figure 4D.
