## Supplementary Figure 6 for "Complex Human hear bearing Skin Organoids reveal Cell Type Specific Susceptibility and Innate Immune Responses to Herpes Simplex Virus 1"

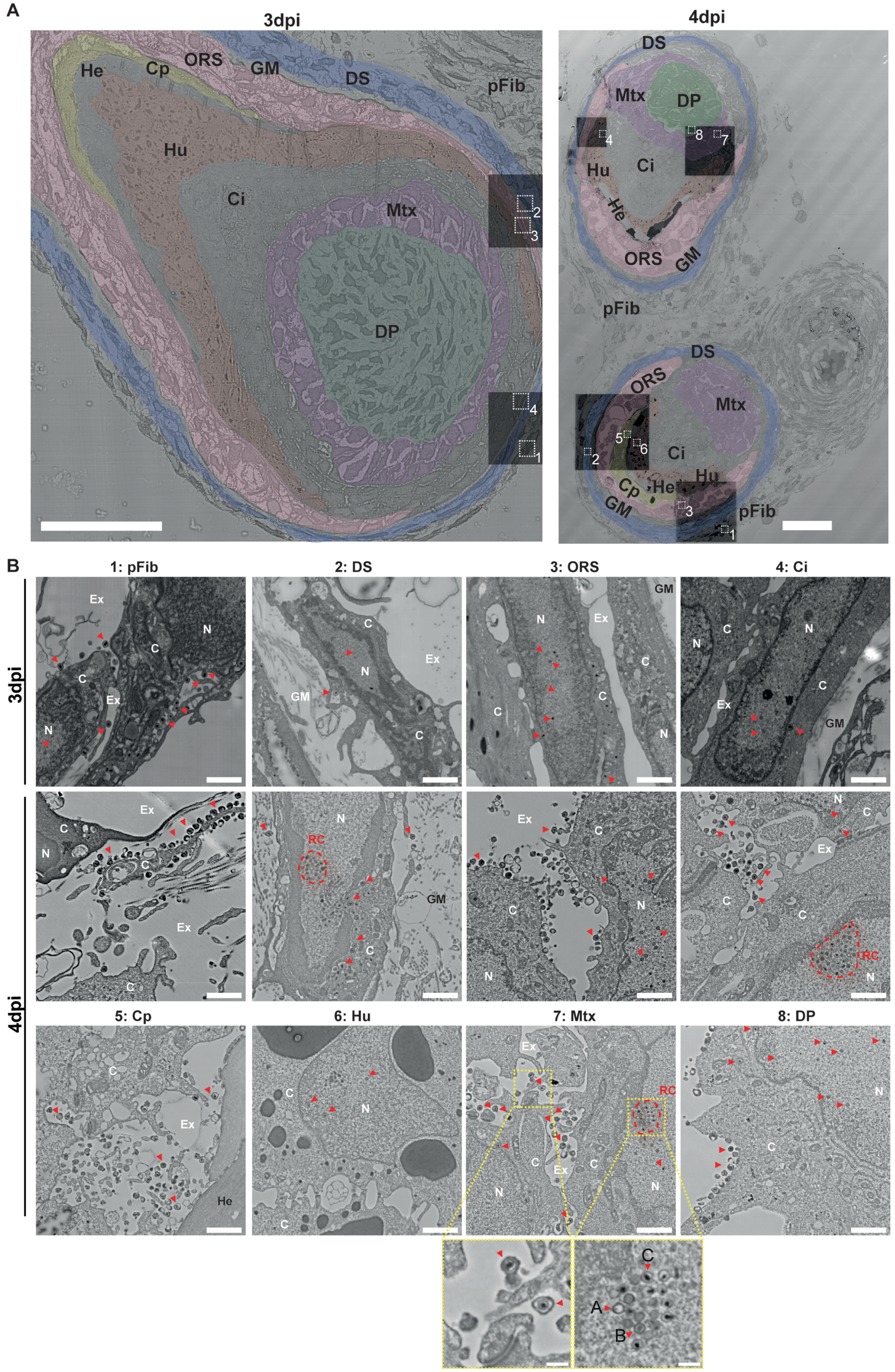


**Supplementary Figure 6: Scanning electron microscopy (SEM) of HFs from HSV-1 infected SkOs at 3dpi and 4 dpi.** **A**: Overviews showing the scanned areas. Black boxes indicate areas scanned with higher resolution. White boxes with numbers indicate areas of zoom-ins shown in B. pFib: papillary fibroblasts; DS: dermal sheath; GM: glassy membrane; ORS: outer root sheath; parts of inner root sheath (IRS): Cp: companion layer, He: Henle's layer, Hu: Huxley's layer, Ci: IRS cuticle; Mtx: matrix; DP: dermal papilla. Scale bar: 50µm. **B:** Zoom-ins on areas indicated in part A. Red arrows indicate viral particles. Yellow boxes indicate areas of zoom-ins showing extracellular viral particles and a nuclear replication compartment with A, B and C capsids. RC: viral replication compartment; Ex: extracellular space, N: nucleus, C: cytoplasm. Scale bar: 1µm; 200nm in zoom-ins.
