## Supplementary Figure 7 for "Complex Human hear bearing Skin Organoids reveal Cell Type Specific Susceptibility and Innate Immune Responses to Herpes Simplex Virus 1"

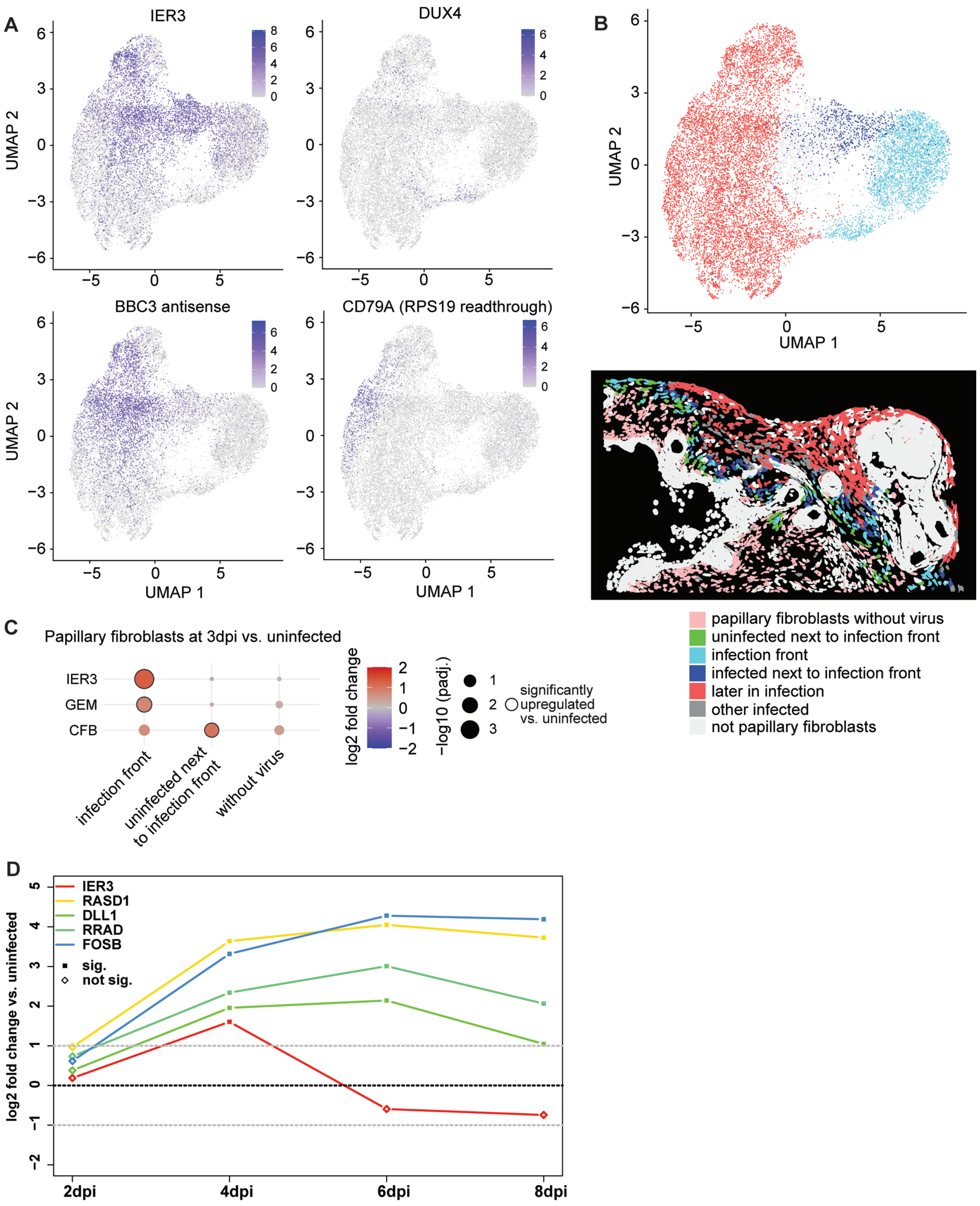


**Supplementary Figure 7: A:** Two-dimensional projection of papillary fibroblasts as in Figure 5A, with cells colored by expression levels of the indicated host genome transcripts. **B:** Papillary fibroblasts were categorized in "infection front" (cluster 1 in Fig. 5AB), "uninfected next to infection front", "infected next to infection front", "later in infection" and "no virus" (i.e., without viral RNA and further away from the infection front). Top: two-dimensional projection as in A, bottom: the same region as in Figure 5B, both with cells colored by these categories. **C:** Differential gene expression in papillary fibroblasts in the cell categories shown in B at 3dpi vs uninfected. Shown are the genes with padj < 0.05 in at least one category. **D:** Fold changes in the bulk RNA-seq data at the indicated timepoints vs. uninfected organoids for IER3 and selected induced host genes. Significantly upregulated genes have padj <0.05.
