## Supplementary Figure 8 for "Complex Human hear bearing Skin Organoids reveal Cell Type Specific Susceptibility and Innate Immune Responses to Herpes Simplex Virus 1"

A

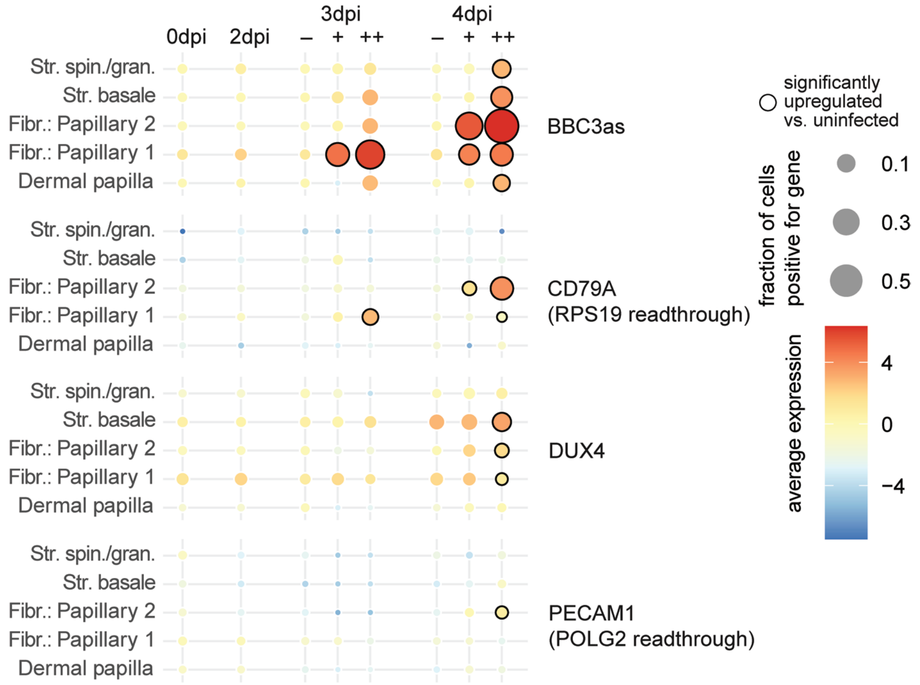


B

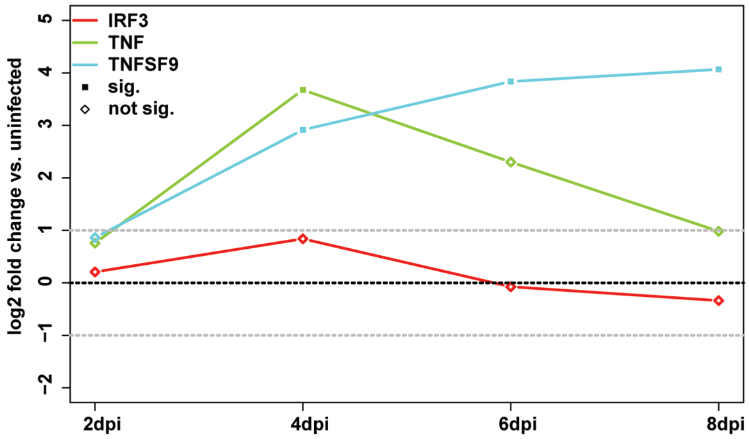


C

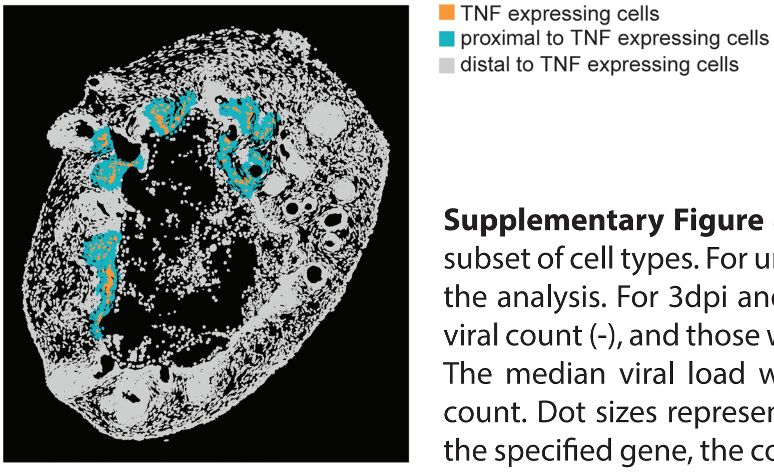


**Supplementary Figure 8: A:** Expression values of the indicated genes in a subset of cell types. For uninfected and 2dpi SkOs, all cells were combined for the analysis. For 3dpi and 4dpi, cells are split up in those with no or only 1 viral count (-), and those with viral load below (+) and above (++) the median. The median viral load was calculated only within cells with more than 1 count. Dot sizes represent the fraction of cells having at least one count of the specified gene, the color the average expression. Black circles around the dots indicate significant differential expression in a pseudobulk comparison vs. all cells in uninfected cells (padj <0.05). **B:** Fold changes in the bulk RNA-seq data at the indicated timepoints vs. uninfected organoids for IRF3, TNF and TNFSF9. Significantly upregulated genes have padj <0.05. **C:** Shown is one replicate of a HSV-1 infected SkO at 4dpi (top left of the 4dpi organoids in Supplementary Figure 3A and Figure 7C). Cells are colored by TNF expressing cells, proximal and distal cells as indicated.
