## Supplementary Table 1 for "Complex Human hear bearing Skin Organoids reveal Cell Type Specific Susceptibility and Innate Immune Responses to Herpes Simplex Virus 1"

**Supplementary Table 1:** Custom panel CZBXK3 for Xenium Spatial transcriptomics

| | | |
|---|---|---|
| ACAN | KLF18 | TP63 |
| AGA | KRT10 | TRIM43 |
| ANKRD1 | KRT14 | TSLP |
| AQP1 | KRT15 | TUBB3 |
| ATOH1 | KRT17 | UCHL1 |
| BBC3 | KRT5 | UTP6 |
| BBC3as | KRT71 | VIM |
| CAMP | LHX2 | ZSCAN4 |
| CD207 | LOR | |
| CD38 | MAVS | |
| COL23A1 | MPZ | |
| COL2A1 | MX1 | |
| DDX58 | MX2 | |
| DEFB103A | NCAM1 | |
| DUX4 | NECTIN1 | |
| EDAR | NEFH | |
| GATA3 | NFATC1 | |
| HSV1_LAT | NFKBIA | |
| HSV1_UL27 | NPNT | |
| HSV1_UL29 | PLP1 | |
| HSV1_UL54 | PMEL | |
| HSV1_US1 | PRAMEF13 | |
| IDO1 | PRPH | |
| IER3 | RNASE7 | |
| IFI44 | RORC | |
| IFNA1 | RSAD1 | |
| IFNAR1 | RSAD2 | |
| IFNB1 | S100B | |
| IFNG | SARM1 | |
| IFNGR1 | SCD | |
| IFNGR2 | SLC27A4as | |
| IL13 | SLC9A1 | |
| IL17C | SOX10 | |
| IL17RE | SRSF3rt | |
| IL22 | SRSF6rt | |
| IL25 | STAT1 | |
| IL4 | STAT2 | |
| IRF3 | TBK1 | |
| IRF7 | TBX21 | |
| IRF9 | TLR3 | |
| ISL1 | TMEM173 | |
| ITGA6 | TNF | |
| ITGA8 | TNFRSF14 | |
| ITGAM | TNFSF9 | |
| ITGAX | TP53 | |
