## Supplementary Table 2 for "Complex Human hear bearing Skin Organoids reveal Cell Type Specific Susceptibility and Innate Immune Responses to Herpes Simplex Virus 1"

**Supplementary Table 2:** Cell type annotation of cell clusters from main and pilot spatial transcriptomics experiment

| Cell type | Key markers (positive) | Key markers (negative, if relevant) | Spatial localization | Notes | Cluster # main experiment | Cluster # pilot experiment |
|---|---|---|---|---|---|---|
| Adipocytes/sebaceous glands | ADIPOQ, PLIN4, LPL, PPP1R1B | | large clusters of cells in dermis and associated with hair follicles | Adipocytes and sebaccous glands are not dinstinguishable based on markers alone, only by localization | 23 | 38 |
| Arrector pili muscle | MEF2C, EDNRB, MYH11, ITGA8,MYLK, ACTA2 | | cells in strands, associated with hair follicles | | 30 | 16 |
| Chondrocytes | ACAN, COL2A1 | | tail | | 4; 36 | 7; 55 |
| Ciliated epithelial cells | CLIC6, SCB2A1, C1orf194,FOXJ1, PCB4, DNAAF1, CCDC39, C6orf118, CFAP53 | | single structure with lumen in one HSV-1 infected SkO at 3dpi | probably an off-target differentiation | 37 | not found |
| Endothelial cells | CD34, CD93, PECAM1, LYVE | | strands in dermis and at outer rim | | 20 | 32 |
| Fibroblasts: Dermal sapilla | SOX2, SOX18, CXCR4 | | base of hair follicle | | 7 | 10 |
| Fibroblasts: Dermal sheath | MFAP5, ACTA2 | | around hair follicle | | 8 | 4 |
| Fibroblasts: Dermal sheath (cyc) | MFAP5, ACTA2, TOP2A, CDK1, MKI67, CENPF, UBE2C | | around hair follicle | cycling dermal sheath cells | 24 | not found |
| Fibroblasts: Mesenchymal type 1 | ASPN, GPC3 | SFRP4 | cells in dermis assoiated with chondrocytes | | 0 | 5 |
| Fibroblasts: Mesenchymal type 2 | ASPN, GPC3, SFRP4 | | single cells in dermis | | 33 | not found |
| Fibroblasts: Myofibroblasts | DES, ASPN, SFRP2, MYBPC1, NCAM1 | | single cells in dermis | | 16; 41 | 25 |

| | | | | | | |
|---|---|---|---|---|---|---|
| Fibroblasts: others | GPC3, PRG4, C7, TMEM100 | | single cells in dermis | | 34 | 40 |
| Fibroblasts: Papillary type 1 | APCDD1, PTGDS, GPC3, PDGFRA, POLCE | | outer rim of dermis | Cells of clusters 5 and 11 (main exp.) showed similar localization as cluster 2 (main exp.). High viral gene expresssion and downregulation of fibroblast markers led to assignment in different clusters. | 2; 5; 11 | 0 |
| Fibroblasts: Papillary type 2 | APCDD1(high), PTGDS, GPC3, PDGFRA, POLCE, MEF2C | | in dermis below epidermis, partially associated with hair follicles | Cells of clusters 13 (main exp.) and 12 (pilot exp.) showed similar localization as clusters 1 (main exp.) and 2 (pilot exp.). High viral gene expresssion and downregulation of fibroblast markers led to assignment in different clusters. | 1;13 | 2;12 |
| Fibroblasts: Papillary type 3 | APCDD1, GPC3, PDGFRA | THBS2, PTGDS, POLCE | single cells in dermis | | 6 | not found |
| Fibroblasts: Pro-inflammatory | PDGFRA, TNC, UCHL1, PDPN, IRF9, STAT2, IFNAR1 | | single cells in dermis | | 12 | not found |
| Keratinocytes: Bulge region | LHX2, LGR5, KRT5, KRT15, NFATC1 | | outer layer at hair follicles | | 10 | 8 |
| Keratinocytes: IRS | KRT71 | | inner layer at hair follicles | | 14 | 14 |

| | | | | | | |
|---|---|---|---|---|---|---|
| Keratinocytes: Matrix | LHX2, LRG5, TOP2A, CDK1, MKI67, CENPF, UBE2C | | hair follicle bulb | | 21 | 17 |
| Keratinocytes: ORS/companion layer | KRT15, TP63, SERPINB3, C5orf46, C15orf48, FGFBP1, SEMA3C, PRDM1 | | outer layer at hair follicle | | 17 | 18 |
| Keratinocytes: Stratum basale | COL17A1, KRT5, KRT15 | | basal layer in epidermis | | 3 | 1 |
| Keratinocytes: Stratum corneum | LOR, SERPINB2, SERPINB3, KRT10, KRT14 | COL17A1, KRT5, KRT15 | higher layers in epidermis | | 18; 25 | 6 |
| Keratinocytes: Stratum spinosum/granulosum | COL17A1, KRT5, KRT15, SERPINB2, SERPINB3, KLK11, KRT10, LOR | | above stratum basale in epidermis | | 9 | 11; 47 |
| Melanocytes | MLANA, PMEL, KIT, SOX10 | | single cells in basal layer and at hair follicles | | 15 | 22 |
| Merkel cells | ATOH1, HEPACAM2, KRT17, CHGA | | single cells at hair follicles | | 43 | 37 |
| Neuronal cells type 1 | TUBB3, SOX2 | | tail of HSV-1 infected SkO at 4dpi | | 19 | not found |
| Neuronal cells type 2 | UCHL1, PTN, MST, SOX2, TUBB3, NCAM1, CHGA, MCF2L, SH2D3C | | tail of HSV-1 infected SkO at 4dpi | | 38 | not found |
| Schwann cells | SOX10, MPZ, S100B | | covering potential axons | | 22 | not found |
| Undefined | too few markers for identification | | dying cells in the middle of the head and highly infected cells | Clusters contain dying cells in the middle of the head and highly infected cells with little to no host gene expression. A cell type assignment was not possible. | 26-29; 31; 32; 35; 39; 40; 42; 44-47 | 3; 9; 13; 15; 19-21; 23; 24; 26-31; 33-36; 39; 41-46; 48-54; 56 |
