## Supplementary Table 3 for "Complex Human hear bearing Skin Organoids reveal Cell Type Specific Susceptibility and Innate Immune Responses to Herpes Simplex Virus 1"

**Supplementary Table 3:** Antibodies used for whole mount immunostaining (WMI) and immunohistochemistry (IHC)

| antibody | vendor | catalog # | dilution WMI | dilution IHC |
|---|---|---|---|---|
| GFP | Abcam | ab290 | - | 1:1000 |
| HSV-1 gC | Santa Cruz | sc-51626 | 1:50 | - |
| HSV-1 gD | Santa Cruz | sc-21719 | - | 1:50 |
| HSV-1 ICP8 | Abcam | ab20194 | 1:50 | - |
| IER3 | Abcam | ab65152 | 1:75 | - |
| IRF3 | Cell Signaling | #11904 | 1:200 | - |
| Ki67 | BD Biosciences | 550609 | 1:100 | 1:100 |
| KRT 15 | Santa Cruz | sc-47697 | 1:50 | 1:100 |
| KRT 17 | Santa Cruz | sc-393091 | 1:100 | 1:200 |
| KRT 20 | Cell Signaling | 13063S | 1:100 | 1:100 |
| KRT 71 | Invitrogen | PA5-83965 | 1:500 | - |
| LHX2 | Millipore | ABE1402 | 1:1000 | 1:750 |
| NEFH | Cell Signaling | 2836S | 1:100 | 1:100 |
| PDGFRα | Cell Signaling | 5241 | 1:100 | 1:200 |
| S100β | Abcam | ab52642 | 1:200 | 1:200 |
| SCD1 | Sigma-Aldrich | HPA012107 | - | 1:50 |
| SOX2 | BD Biosciences | 561469 | 1:100 | 1:100 |
| TUBB3 | BioLegend | 801202 | 1:100 | 1:100 |
