## Supplementary Note for "Complex Human hear bearing Skin Organoids reveal Cell Type Specific Susceptibility and Innate Immune Responses to Herpes Simplex Virus 1"

### Processing and cell type annotation of spatial transcriptomics data

For the main experiment, three organoids per timepoint (uninfected as well as 2/3/4dpi) were measured on a Xenium slide. We used the pre-designed human multi-tissue and cancer panel with 377 genes as well as a custom 100 gene panel including 5 viral genes (supplementary table 2) for multiplexed transcript detection. For improved cell segmentation, we used the cell segmentation add-on kit, which allows segmentation based on DAPI staining, internal protein and rRNA detection, as well as cell surface staining. Raw data including cell segmentation was pre-processed on the Xenium device using manufacturer's defaults to yield the initial data folder, available through NCBI GEO (accession number GSE313919).

Subsequently, data was read into R for further analysis. Very low quality cells (less than 10 counts) were filtered out. To increase sensitivity for cell type detection, we included three uninfected organoids from an infection experiment run in parallel using skin organoids and Merkel cell polyoma virus infection[1]. Next, cells from all time points (i.e., three organoids each from this experiment uninfected/2/3/4dpi as well as three uninfected organoids from the polyoma virus experiment, in total 15 skin organoid sections) together were clustered based on their transcriptome using Seurat[2] (Supplementary Note Figure 1). Expression of marker genes across clusters was used to manually assign clusters to cell types (Supplementary Note Figure 2, Supplementary Table 3). Next to major fibroblast and keratinocyte clusters, we found a range of less abundant cell types (Supplementary Note Figure 3). Some clusters were assigned as "undefined" cells, which frequently contained high amounts of viral RNAs (Supplementary Note Figure 4). Likely, as a consequence cell type identity could not be assigned due to loss of marker genes. Of note, we also attempted cell type annotation following integration using harmony[3] or SCtransform[4], as well as annotation using the BuildNicheAssay function in the Seurat package[2], or BANKSY[5]. While these algorithms successfully identified the expected major cell types, they failed to detect low-abundance populations such as melanocytes and Merkel cells, which occur as single cells embedded within clusters of another cell type, for example in the stratum basale. In contrast, cell-type annotation based solely on cellular transcriptomes was superior, enabling robust detection of both major cell types and these rare populations.

### Pilot experiment

Prior to the main experiment described above, which is the one presented in the main figures, we performed a pilot experiment. We used the same gene panels as described above, however with the timepoints uninfected/2/6dpi, and with DAPI-staining based nuclear expansion for cell segmentation instead of the cell segmentation add-on kit. Cells were re-segmented with a 10 µm expansion distance instead of the standard 15 µm distance using CellRanger (10x Genomics). Otherwise, cell types were assigned as described above for the main experiment (Supplementary Note Figures 5 and 6). In contrast to the main experiment, we failed to identify several low-abundant cell types, which could be due to decreased sensitivity with the nuclear expansion method. At 6dpi, most cells could not be assigned to a cell type, likely due to high amounts of viral RNA across the entire organoid sections and the virus induced host cell shutoff. From the pilot experiment, we concluded that advanced cell segmentation using the cell segmentation add-on kit which we used in the main experiment was superior to nuclear expansion–based cell type annotation. In addition, by 6dpi, HSV-1 infection had progressed too far to be suitable for spatial transcriptomics experiments.


1 Albertini, S. *et al.* Merkel cell polyomavirus infection and persistence modelled in skin organoids. *bioRxiv*, 2025.2002.2011.637697 (2025). https://doi.org:10.1101/2025.02.11.637697

2 Hao, Y. *et al.* Dictionary learning for integrative, multimodal and scalable single-cell analysis. *Nat Biotechnol* **42**, 293-304 (2024). https://doi.org:10.1038/s41587-023-01767-y

3 Korsunsky, I. *et al.* Fast, sensitive and accurate integration of single-cell data with Harmony. *Nat Methods* **16**, 1289-1296 (2019). https://doi.org:10.1038/s41592-019-0619-0

4 Hafemeister, C. & Satija, R. Normalization and variance stabilization of single-cell RNA-seq data using regularized negative binomial regression. *Genome Biol* **20**, 296 (2019). https://doi.org:10.1186/s13059-019-1874-1

5 Singhal, V. *et al.* BANKSY unifies cell typing and tissue domain segmentation for scalable spatial omics data analysis. *Nat Genet* **56**, 431-441 (2024). https://doi.org:10.1038/s41588-024-01664-3

**Supplementary Note Figure 1:** UMAP representation of spatial transcriptomics data from main experiment

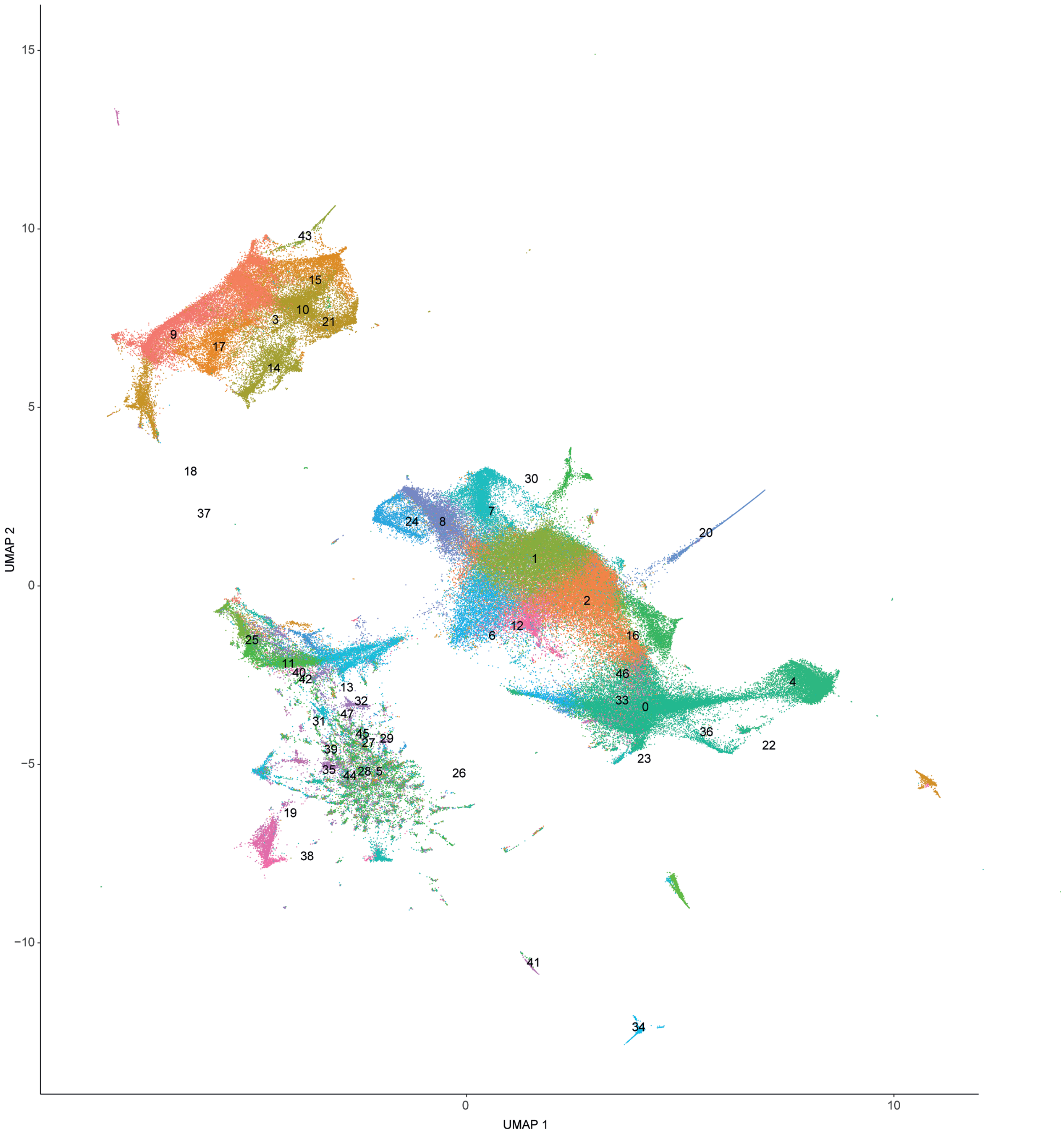

**Supplementary Note Figure 2:** Expression of marker genes in clusters from main experiment

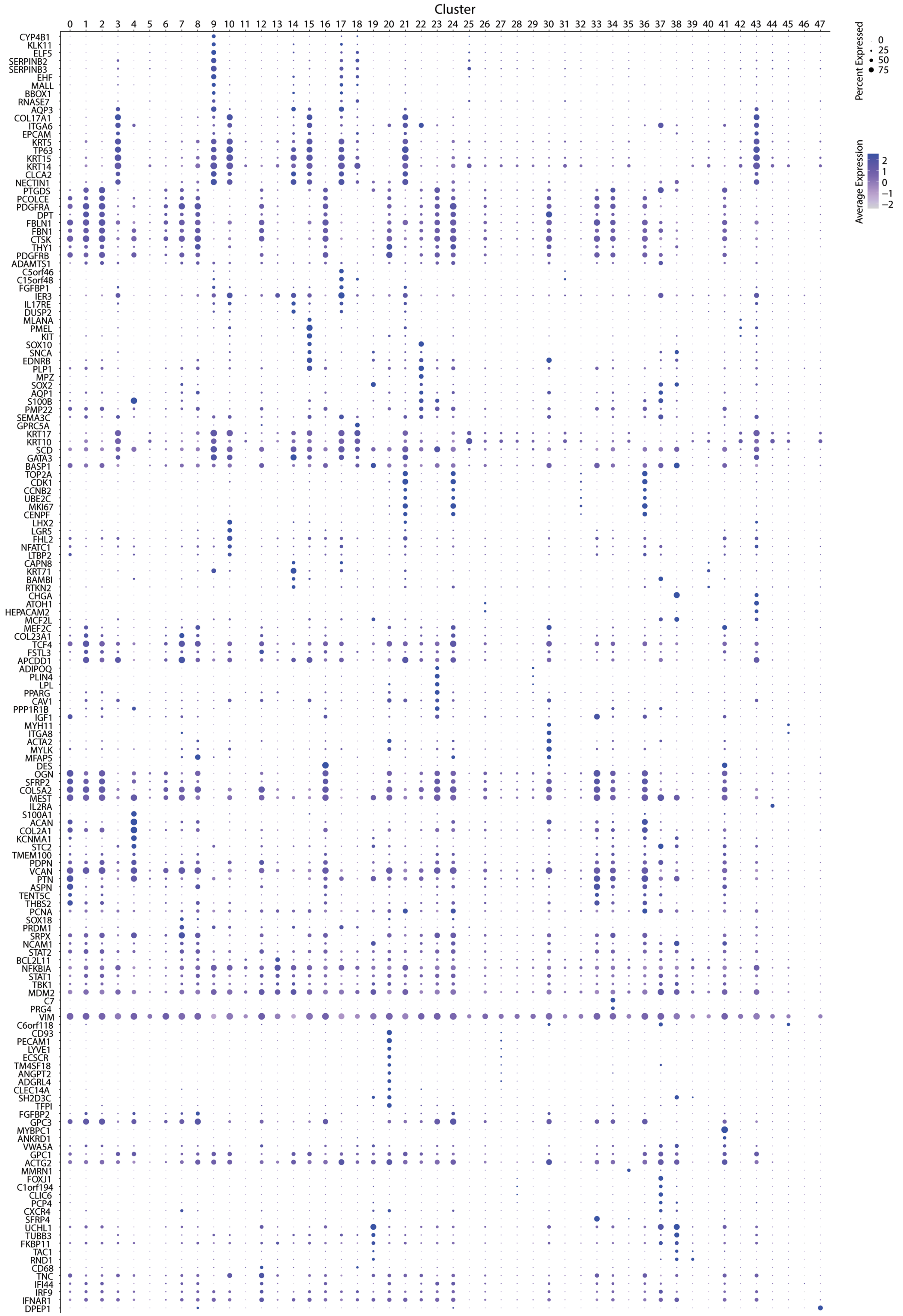

**Supplementary Note Figure 3:** UMAP representation of spatial transcriptomics data from main experiment with annotation of cell types

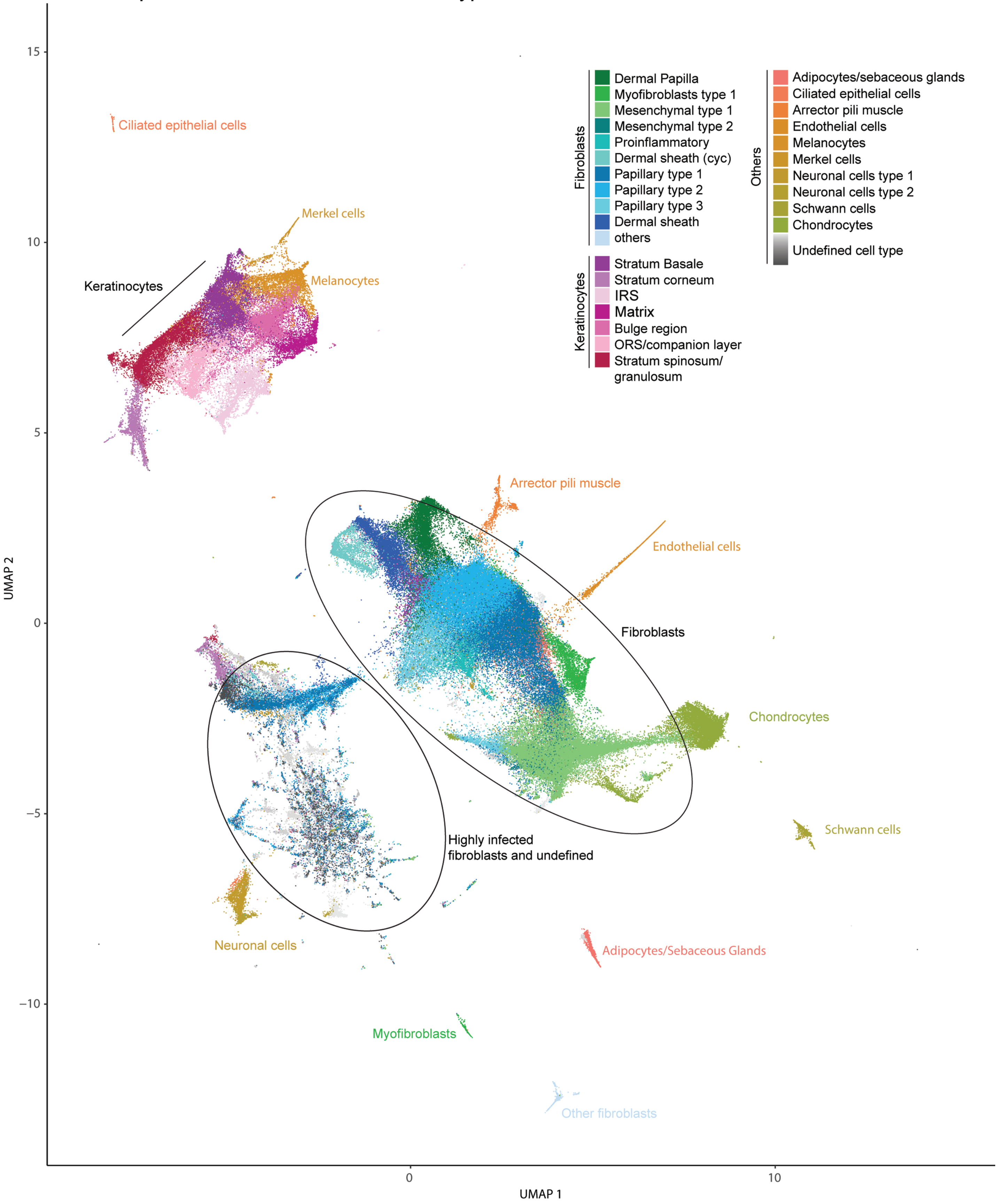

**Supplementary Note Figure 4:** UMAP representation of spatial transcriptomics data from main experiment colored by normalized HSV-1 UL54 expression

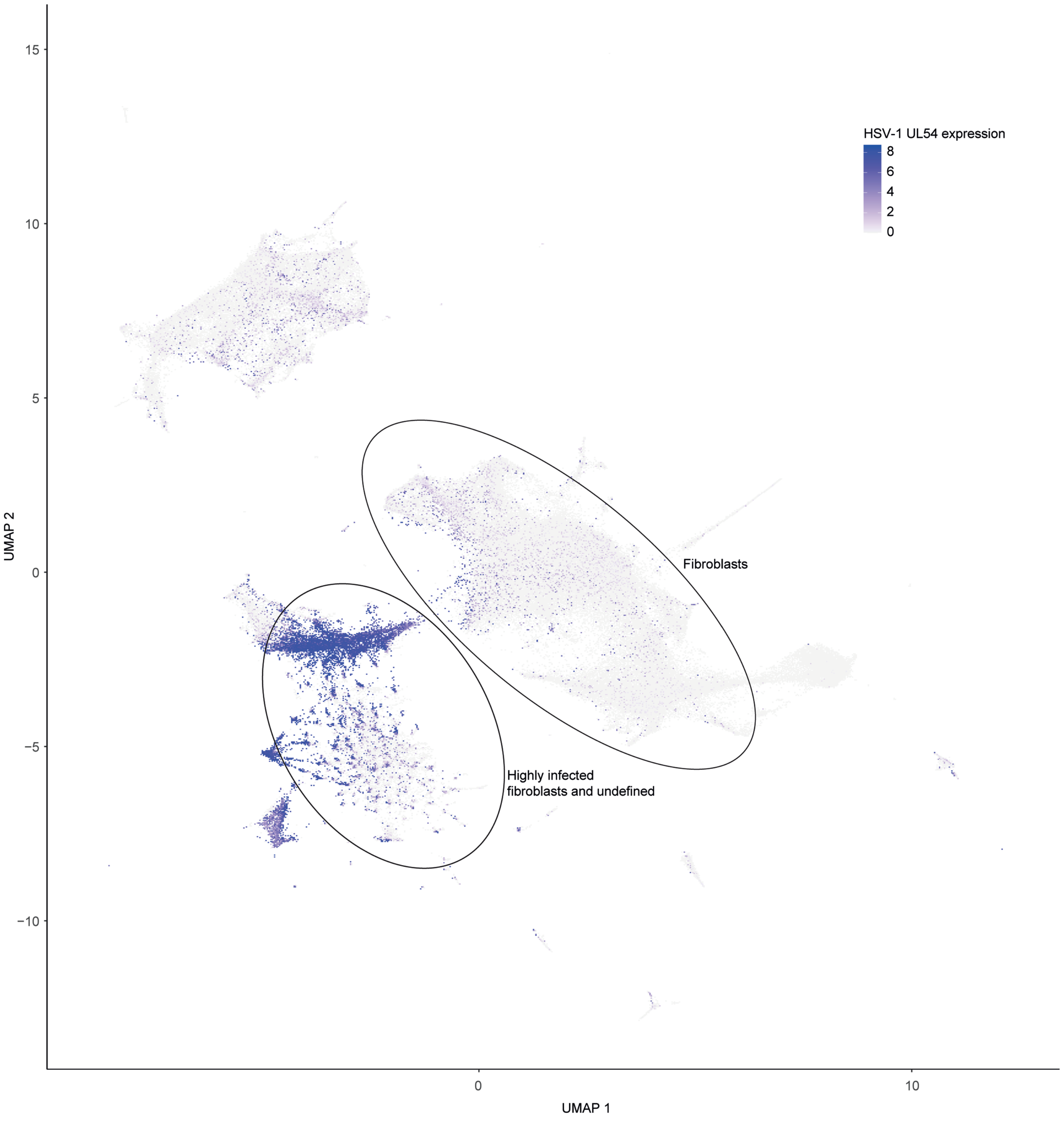

**Supplementary Note Figure 5:** Overview of all SkOs analyzed by spatial transcriptmics in the pilot experiment with cells colored by cell type

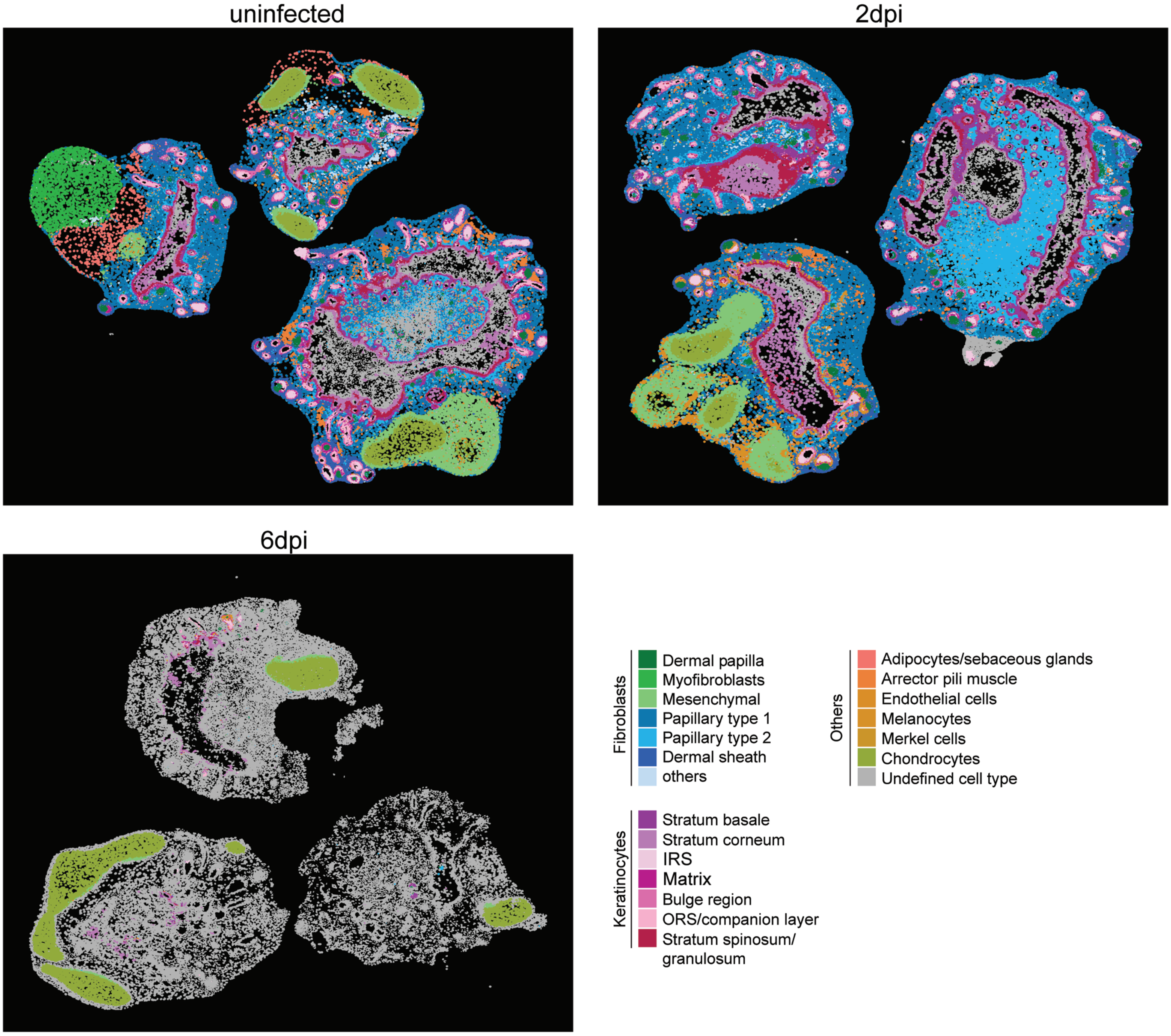

**Supplementary Note Figure 6:** Overview of all SkOs analyzed by spatial transcriptomics in the pilot experiment with cells colored by normalized HSV-1 UL54 expression

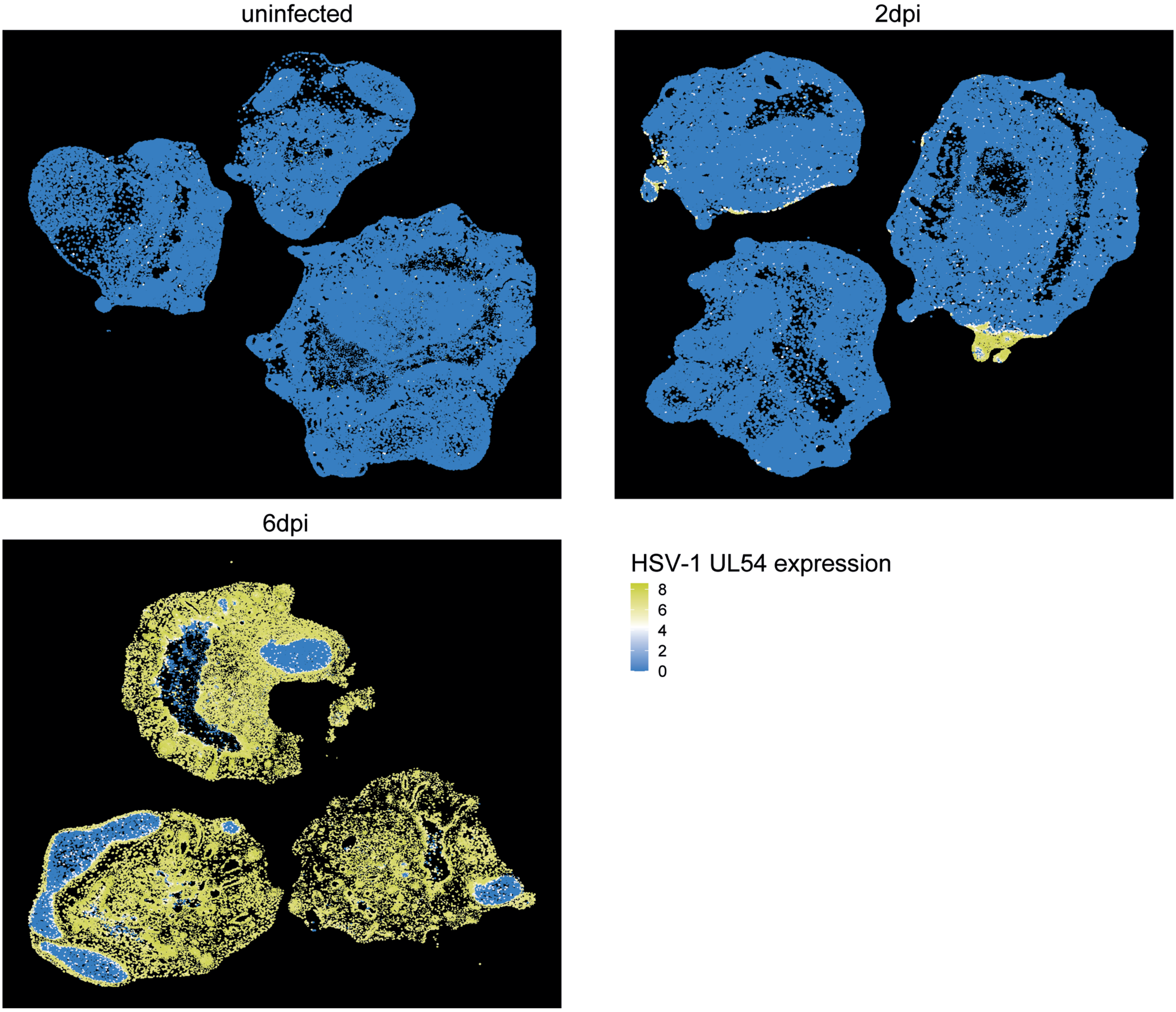

**Supplementary Table 1:** Custom panel CZBXK3 for Xenium Spatial transcriptomics
